# Genetic diversity and evidence of recombination of *Horsegram yellow mosaic virus* infecting pole bean (*Phaseolus vulgaris* L.) from South India

**DOI:** 10.1101/2023.03.27.534342

**Authors:** J Bindu, H. D. Vinay Kumar, Shridhar Hiremath, Mantesh Muttappagol, M. Nandan, Devaraj, C. R. Jahir Basha, K. S. Shankarappa, V. Venkataravanappa, C. N. Lakshminarayana Reddy

## Abstract

The yellow mosaic disease (YMD) caused by begomoviruses is a major constraint for the production of pole bean (*Phaseolus vulgaris* L.) in India. Survey was carried out in the eastern dry zone of Karnataka during 2019-20 to record the incidence of yellow mosaic disease in pole bean which revealed the ubiquitous prevalence of YMD in pole bean ranging from 6.02 to 80.74 per cent. Leaf samples collected (symptomatic and asymptomatic) were subjected for begomovirus detection using specific primers. Twelve samples, representing all the 12 taluks in the surveyed region were considered for full genome amplification by RCA, cloned and sequenced. Genome length of 12 current isolates ranged from 2718 – 2744 and 2668 – 2671 nucleotides for DNA-A and DNA-B, respectively. Sequence analysis using Sequence Demarcation Tool (SDT) showed >91 per cent nucleotide identity of current isolates (DNA-A) with other horsegram yellow mosaic virus (HgYMV) isolates available in the GenBank. As per existing ICTV criteria, all the current isolates can be considered as strains of HgYMV. Further, DNA-B associated with all the 12 isolates also shared >91 per cent nucleotide identity with DNA-B of HgYMV isolates, indicating absence of component re-assortment in HgYMV. Variation in the pairwise nucleotide identity and phylogenetic analysis confirmed the existence of new strains within the current HgYMV isolates. GC plot analysis reveals potential recombination in the low GC rich regions. Further, recombination breakpoint analysis indicated intra-species recombination in both DNA-A and DNA-B, which might have driven the origin of new strains in HgYMV. This is the first comprehensive study on begomoviruses ioslates associated with the yellow mosaic disease of pole bean based on complete genome sequencing in the world.

## 1 Introduction

Begomoviruses have single-stranded (ss) DNA genome encapsidated in twinned icosahedral particles, belongs to the family, *Geminiviridae* and comprises 14 genera, which are classified based on genome structure, host range and insect vectors (Zerbini et al., 2017). Among the 14 genera, the most destructive and widespread viruses belong to the genus, Begomovirus, which are transmitted by whitefly, *Bemisia tabaci* (De Barro et al., 2011). Based on their genome organization, begomoviruses have been divided into monopartite (having homologue of DNA-A bipartite viruses) or bipartite (both DNA-A and DNA-B components) (Hanley-Bowdoin et al., 2013). Genomes of the monopartite and bipartite begomoviruses encode proteins that are required for transmission and replication of the genome. The DNA-A genome has six open reading frames (ORFs) which codes for different proteins *viz.,* replication associated protein (AC1) which initiates the replication of viral DNA, transcription activator protein (AC2) helps in transactivation of expression of negative sense gene expression, replication enhancer protein (AC3) play a vital role in efficient replication of viral DNA, AC4 protein involved in virus infectivity and coat protein (CP) coded by AV1 and AV2 involved in encapsidation and insect transmission (Hanley-Bowdoin et al., 2013). DNA-B genome has two ORFs *viz*., BC1 which encodes for movement protein (MP) for cell to cell movement and BV1 which encodes for the nuclear shuttle protein (NSP) responsible for suppression of transmembrane receptor kinase activity, both of which are vital in systemic spread and symptom development (Brown et al., 2015). DNA-A and DNA-B share a common region (CR) of approximately 200 nucleotides (nt) long which is highly conserved in both the components. This region contains repeated motifs (iterons), which are having sequence-specific replicase-binding sites and a stem-loop structure with the nonanucleotide (TAATATTAC) in the loop, conserved feature among geminiviruses and known to play a crucial role in replication of the viral genome (Fauquet et al., 2003, Martín-Hernández and Pagan, 2022).

Pole bean (*Phaseolus vulgaris* L.), a member of the *Fabaceae* (legume) family, is a predominantly self-pollinated crop, grown worldwide for its edible green pods and dry seeds. Pole bean has recognized benefits to human health and nutrition, as it is rich source of protein (22%), carbohydrate (62%), soluble fibers (15%) and micronutrients like calcium, iron, magnesium, phosphorus and potassium (Jeevan et al., 2015). Dry seeds and fresh green pods are used as vegetables in the diet, predominantly among the vegetarian population of India. In India, pole bean is grown in an area of 227.78 thousand hectares with the production of 2276.95 thousand metric tonnes and productivity of 9 tonnes per hectare (https://www.indiastat.com/).

It is prone to various diseases like angular leaf spot, bacterial brown spot, common blight, rust, root rot, anthracnose (Nene et al., 1988). Apart from these, pole bean succumbs to different viral diseases namely, bean golden mosaic, common bean mosaic, bean yellow mosaic, bean leaf roll, tobacco curly shoot, bean pod mottle (Nene, 1978).

In southern Asia, grain legume crops are suffering huge loss due to attack of different begomoviruses (Ilyas et al., 2009; Qazi et al., 2007). Begomoviruses infecting legumes are grouped under legumoviruses, which mainly consist of horsegram yellow mosaic virus (HgYMV), dolichos yellow mosaic virus (DoYMV), mungbean yellow mosaic virus (MYMV), mungbean yellow mosaic India virus (MYMIV), which are known to cause yellow mosaic disease (YMD) of leguminous crop in India (Agnihotri et al., 2019). Apart from these, other viruses which are reported on legume crops are tobacco curly shoot virus (TbCSV) (Venkataravanappa et al., 2012), french bean leaf curl virus (FbLCV) (Kamaal et al., 2013), tomato leaf curl Gujarat virus (ToLCGV) (Kamaal et al., 2015) and tomato leaf curl Joydebpur virus (ToLCJV) (Ansar et al., 2019). In recent years, cultivation of pole bean in India has suffered serious losses due to yellow mosaic disease (YMD) (Archith et al., 2017). Based on the CP gene sequence analysis, YMD of pole bean was confirmed to be associated with begomovirus infection (Jeevan et al., 2015). However, the exact strains of begomovirus infecting pole bean was not confirmed due to the unavailability of full-length sequence of the begomovirus, which is essential for nomenclature of any begomovirus. With this backdrop, the present study was carried out aiming to determine prevalence of disease and characterize begomovirus associated with YMD of pole bean from India.

## 2 Material and Methods

### 2.1 Field survey and collection of virus-infected pole bean samples

A roving survey was conducted during 2019-20 to assess the per cent disease incidence (PDI) of YMD on pole bean in different farmer’s fields of Bengaluru Rural (Doddaballapura, Hoskote and Nelamangala), Chikkaballapura (Bagepalli, Chintamani, Chikkaballapur and Sidlaghatta) and Kolar (Bangarapete, Kolar, Malur, Mulbagal and Srinivaspura) districts of Karnataka State, India. The PDI was estimated in each field by visual examination by counting the number of infected plants over the healthy plants in a 20 x 20 meter plot. During the survey varied types of symptoms *viz*., mosaic, yellowing, vein clearing, reduced leaf size, downward curl, deformed fruits and pods with immature seeds were observed on the pole bean plants. A total of 26 fields were surveyed which includes 12 in Kolar, seven each in Bangalore Rural and Chikkaballapura districts (Fig. 1). From each field one symptomatic and asymptomatic leaf samples were collected. The infected samples collected from each field were designated as separate virus isolates (PB-1 to 26).

**Figure 1:**
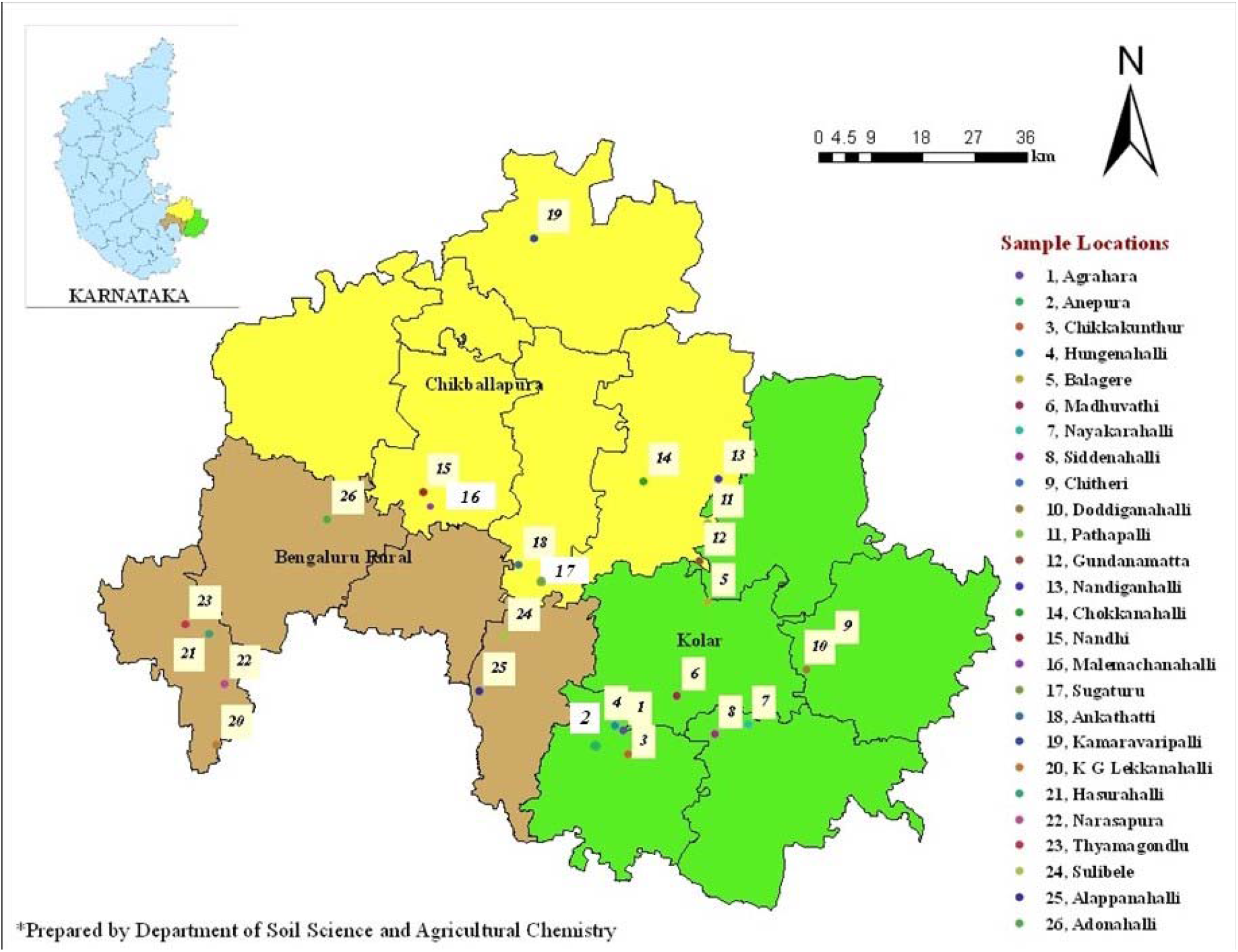
GIS map showing locations surveyed in Kolar, Chikkaballapur and Bengaluru Rural districts of Karnataka state, India for the incidence of YMD of pole bean and plant samples collected (using Arc GIS software).

### 2.2 Nucleic acid isolation, polymerase chain reaction and sequencing

Total genomic DNA was isolated from the symptomatic and asymptomatic pole bean leaf samples using CTAB method (Doyle and Doyle 1990). Initially all 26 samples were subjected to polymerase chain reaction (PCR) using primers specific to the genome of the begomoviruses as described by Venkataravanappa et al., (2012) (Table S1). Partial genome sequencing of all 26 samples indicated that all the samples were associated with begomovirus. Based on phenology symptoms on pole bean and nucleotide (nt) identity, one isolate from each taluk (PB2, PB5, PB8, PB10, PB12, PB13, PB15, PB17, PB19, PB20, PB24 and PB26) was processed for amplification of complete genome of the virus by rolling circle amplification (RCA) method using TempliPhi illustra amplification kit (GE Healthcare, Piscataway, NJ). The resulting RCA product (2 μL) was digested with *Bam*H1 (DNA-A) and *Xba*I (DNA-B) to obtain the monomeric units of the complete genome (2.7kb). The resulting monomeric fragment of the virus was purified using gel extraction kit and ligated into a linearized pUC18 vector as per the manufacturer’s instructions (Fermentas, Germany). The recombinant clones were confirmed by restriction digestion. Three positive clones were selected and the viral genome insert was sequenced by primer walking strategy at Medauxin Pvt. ltd, Bengaluru, India.

### 2.3 Sequence and GC content analysis of bipartite begomovirus genome

The genome sequences (DNA-A and DNA-B) of the begomovirus isolates infecting pole bean obtained in the present study was subjected to Vector NTI Advance TM 9 software (Invitrogen, Foster City, CA, USA) to remove vector sequences. Further, ORFs were identified by using the National Center Biotechnology Information (NCBI) ORF finder tool (http://www.ncbi.nlm.nih.gov/gorf/gorf.html) and their translation was checked by Expasy translation tool (http://www.expasy.org/tools/dna.html). The sequences showing maximum nt similarity with legumoviruses DNA-A (Table S2) and DNA-B (Table S3) were retrieved from the NCBI database. Sequence Demarcation Tool version 1.2 (SDTv1.2) (Muhire et al., 2014) was used to calculate pairwise per cent identity between pole bean isolates of current study with GenBank isolates. With the aid of MEGA X software (Kumar et al., 2018), phylogenetic tree was generated using the Neighbor Joining method applying Kimura 2-parameter with 1000 bootstrap replications. Further, the pole bean isolates were subjected to Artemis DNA plotter analysis tool v18.1.0. (http://www.sanger.ac.uk/Software/Artemis) to generate guanine-cytosine (GC) plot graphs (Euesden et al., 2015). The plots were generated with the parameters such as graph height 0.7, window size 80 and step size 1. The GC-plot graphs help in determining the distribution of GC content throughout the genome. The genomic region with less GC content is considered to be the hot spot region for recombination (Yogindran et al., 2021) and begomoviruses are well known for their recombination potential. To know the possible recombination events in DNA-A and DNA-B components of begomovirus isolates, split-decomposition trees were constructed with 1000 bootstrap replicates in Split Tree version 4.11.3 with default settings (Huson and Bryant, 2006) and recombination breakpoint analysis was carried out using RDP4 with GENECOV, Max Chi, Chimara, Si Scan, 3Seq algorithms (Martin et al., 2015) and 0.05 *p-value* cut off throughout with standard Bonferroni correction.

## 3 Results

### 3.1 Survey for the yellow mosaic disease (YMD) in pole bean and collection of virus infected samples

Survey results revealed that YMD on pole bean is prevalent in all the fields surveyed and PDI ranged from 6.02 to 80.74 per cent. Among three districts surveyed in Karnataka, maximum mean disease incidence was recorded from Kolar district followed by Chikkaballapur and Bengaluru Rural districts with mean disease incidence of 33.91, 33.29, and 28.77 per cent, respectively (Fig. 1 and Table S4). Disease was observed in all growth stages of the crop starting from 15 days after sowing up to the harvesting stage of the crop with typical characteristic symptoms of vein clearing, mosaic, yellowing, reduced leaf size, downward curling, deformed fruits and pods with immature seeds (Fig. 2). If the disease starts at pre-vegetative stage, then yield loss of 100 per cent was observed. Whereas, in case of the disease occurring at reproductive and post reproductive stage led to the loss of yield up to 20 to 25 per cent in all surveyed areas.

**Figure 2:**
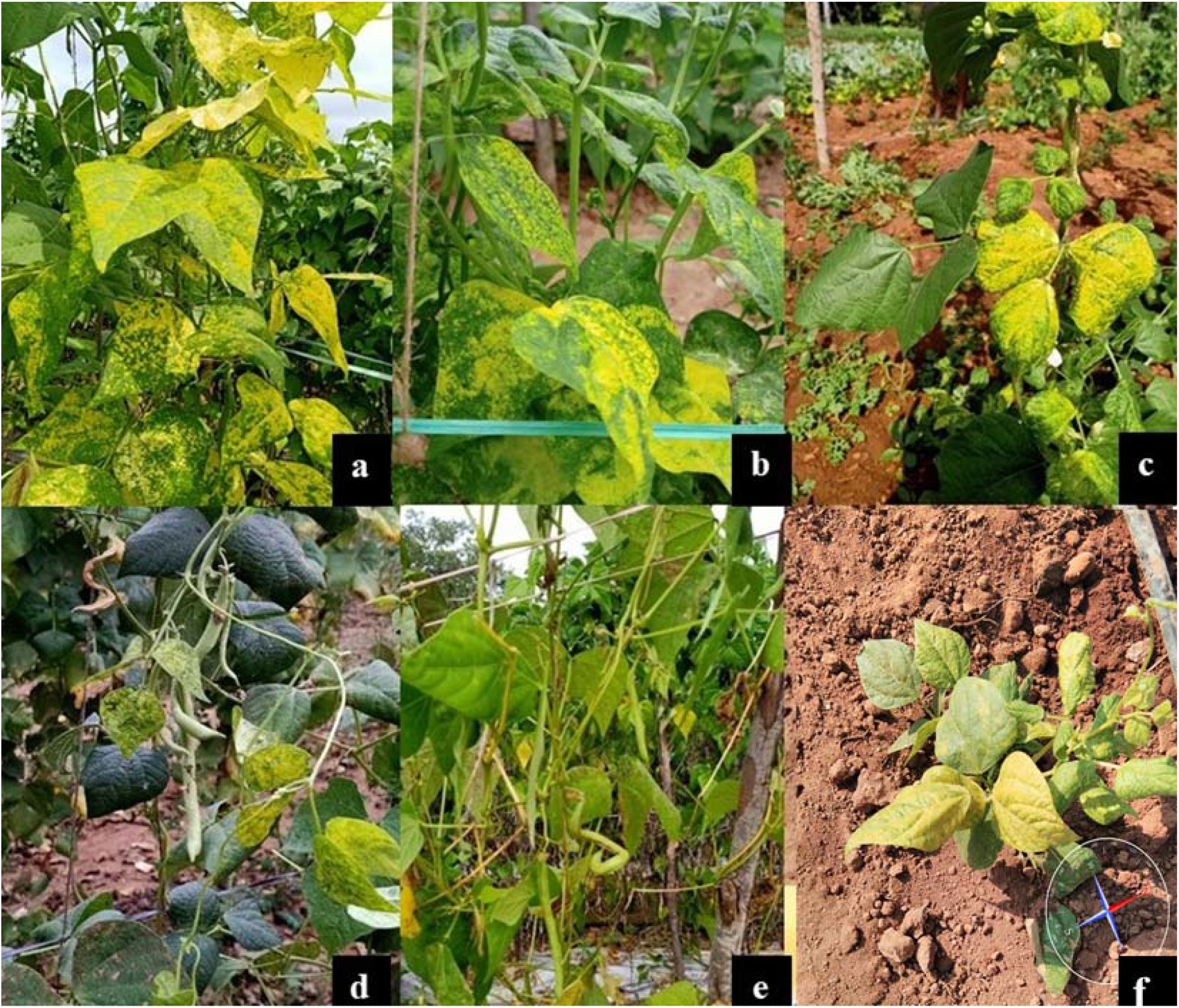
Pictorial representation of symptoms of yellow mosaic disease on pole bean A) and B) Mosaic and complete yellowing C) and D) Vein clearing and downward curling, E) Deformed fruit and F) Stunted growth

### 3.2 Detection of begomovirus in pole bean leaf samples

Symptomatic and asymptomatic pole bean leaf samples collected from different locations of Karnataka, India were confirmed for the presence of begomovirus using specific primers. The partial genome amplification (1.2kb) and sequencing of 26 isolates showed more than 95 per cent nt identity among themselves. Therefore, 12 pole bean isolates were selected for complete genome amplification using the RCA method representing each taluk in the surveyed area. Amplified RCA products of 12 pole bean isolates were cloned and sequenced. Complete nt sequences analysis (DNA-A and DNA-B) of 12 pole bean isolates indicated of having more than 92 per cent nt identity between them and with other HgYMV isolates infecting different crops. Consensus sequences of 12 pole bean isolates sequences are available under accession numbers MW816828-39 (DNA-A) and MW816840-51 (DNA-B) in NCBI database.

### 3.3 Genome structure of DNA-A and DNA-B of begomoviruses associated with pole bean

Complete genome (DNA-A) of 12 pole bean isolates (PB2, PB5, PB8, PB10, PB12, PB13, PB15, PB17, PB19, PB20, PB24 and PB26) ranged from 2718 to 2744 nt in size and exhibits similar genomic structures typical to Old World (OW) bipartite begomoviruses reported so far. The genome encodes for six ORFs in DNA-A molecule; AV2 [Pre-coat protein, 145-495, 351 nucleotide (nt) /116 amino acid (aa)] and AV1 (Coat protein, 305-1078, 774 nt / 257aa) in sense orientation and AC1 (replication-associated protein, 1527-2615, 1089 nt / 369 aa), AC2 (transcriptional activator protein, 1220-1627, 408 nt / 135 aa), AC3 (replicase enhancer protein, 1075-1479, 405 nt/134 aa) and AC4 (C4 protein, 2165-2458, 294 nt /97 aa) in antisense orientation. Similarly, DNA-B molecule encodes two conserved ORFs: BV1 (nuclear shuttle protein, 428-1198, 771 nt/256 aa) and BC1 (movement protein 1233-2129, 879 nt/ 298 aa) and in sense and antisense orientation, respectively.

Complete genome (DNA-A) of 12 pole bean begomovirus isolates in the present study were compared within the collected isolates and sequences of 50 selected begomoviruses infecting different legume crops retrieved from the NCBI database, that includes HgYMV (11), MYMIV (11), MYMV (12), DoYMV (7), velvet bean golden mosaic virus (VbGMV) (4), FbLCV (3), ToLCGV (1) and TbCSV (1). Per cent nt identity within 12 isolates ranged from 95.10 to 98.20. All 12 pole bean isolates (PB2, PB5, PB8, PB10, PB12, PB13, PB15, PB17, PB19, PB20, PB24 and PB26) showed the nt identity with the range of 92.70 to 98.00 per cent with several isolates of HgYMV infecting cowpea, french bean, moth bean, lima bean, horsegram and shared low nt sequence identity (< 86.50%) with several isolates of MYMIV, MYMV, VbGMV, DoYMV, FbLCV, ToLCGV and TbCSV infecting different legumes in India (Table S5). These results were also well supported by the sequence demarcation graph generated using SDTv1.2 (Fig. S1). Based on the current species demarcation criteria for begomoviruses (91% nt sequence identity) (Brown et al., 2015), the begomovirus isolates isolated from pole bean were found to be variants of HgYMV. Phylogenetic analysis revealed that 12 pole bean isolates clustered with several isolates of HgYMV infecting cowpea, french bean, moth bean, lima bean, horsegram crops (Fig.3). Variation in the nt identity of DNA-A component of 12 pole bean isolates and their sub clustering in HgYMV group indicates the presence of strainal variation in the isolates infecting pole bean and hence, current isolates are considered as isolates of HgYMV. The HgYMV isolates in the current study are bifurcated into three strainal groups. First group having 10 HgYMV isolates (PB2, PB5, PB8, PB10, PB12, PB13, PB15, PB17, PB19 and PB20) infecting pole bean clustered separately, second group having one HgYMV isolate (PB24) showed close clustering with HgYMV isolates infecting lima bean and third group with one HgYMV isolate (PB26) clustering with HgYMV isolates infecting french bean and horsegram (Figure 3).

**Figure 3:**
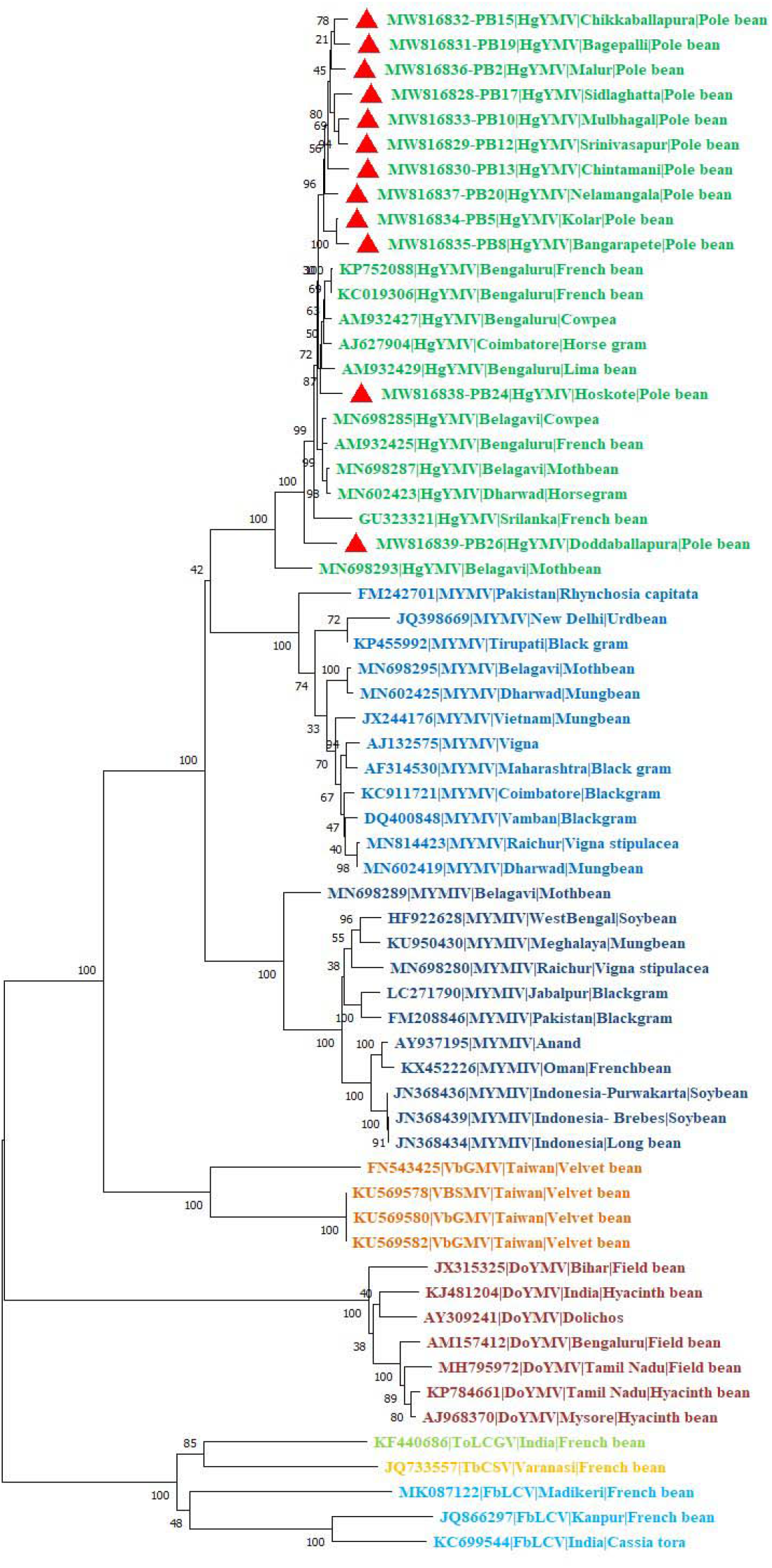
Phylogenetic relationship of nucleotide sequences of DNA-A component of 12 HgYMV isolates from pole bean, with selected sequences of 50 begomoviruses infecting legumes by Neighbour Joining method using MEGA X, with 1000 bootstrap replicates

Comparison of individual ORFs of 12 HgYMV isolates (AV1, AV2, AC1, AC2, AC3 and AC4) from pole bean with other selected sequences of 50 begomoviruses infecting legumes revealed that they shared maximum nt and aa identity with HgYMV isolates and low nt and aa identity with other legume infecting begomoviruses (MYMIV, MYMV, VbGMV, DoYMV, FbLCV, ToLCGV and TbCSV) (Table S5).

The intergenic region (IR) of 12 HgYMV isolates had more homology with IRs of several isolates of HgYMV infecting cowpea, french bean, moth bean, lima bean and horsegram, respectively (Table S5). Length of IR is 264-271 nts and is similar to those of other bipartite begomoviruses reported so far. The IR encompasses an absolutely conserved hairpin structure containing nonanucleotide sequence (TAATATTAC) that marks the origin of virion-strand DNA replication. Two repeated sequences known as “iterons” (GGTGA nt positions 2672-2676 and 2679-2683) were detected adjacent to the TATA box in all 12 HgYMV isolates, which are recognition sequences for binding of the Rep promoter (Hanley-Bowdoin et al., 1999) and share significant homology with iterons identified in DNA-A reported earlier.

Complete nucleotide sequence of DNA-B of 12 HgYMV isolates was compared within collected isolates and other selected sequences of begomoviruses infecting legumes that includes HgYMV (12), MYMIV (13), MYMV (15) and DoYMV (2) retrieved from the NCBI database. Per cent nt identity for DNA B within 12 collected isolates ranged from 94.70 to 97.40. Whereas, 12 HgYMV isolates showed nt identity of 93.70 to 97.40 with several HgYMV isolates infecting cowpea, french bean, moth bean, lima bean, horsegram and low nt identity (< 85%) with other legume infecting begomoviruses (MYMIV, MYMV and DoYMV) (Table S6). This result was well supported by sequence demarcation graph, in which all 12 HgYMV isolates are closely related to several HgYMV isolates infecting different crops (Fig. S2). Further, the phylogenetic analysis of 12 HgYMV isolates from the pole bean were bifurcated into three groups indicates the presence of strainal variation in the HgYMV isolates infecting pole bean. First group having nine HgYMV isolates (PB2, PB5, PB10, PB12, PB13, PB15, PB19, PB24 and PB26) infecting pole bean are clustered separately, second group having one HgYMV isolate (PB17) closely clustering with HgYMV isolates infecting moth bean and mung bean and third group having two HgYMV isolate (PB8 and PB20) segregating into separate cluster with the HgYMV isolates infecting french bean and horsegram (Fig. 4).

**Figure 4:**
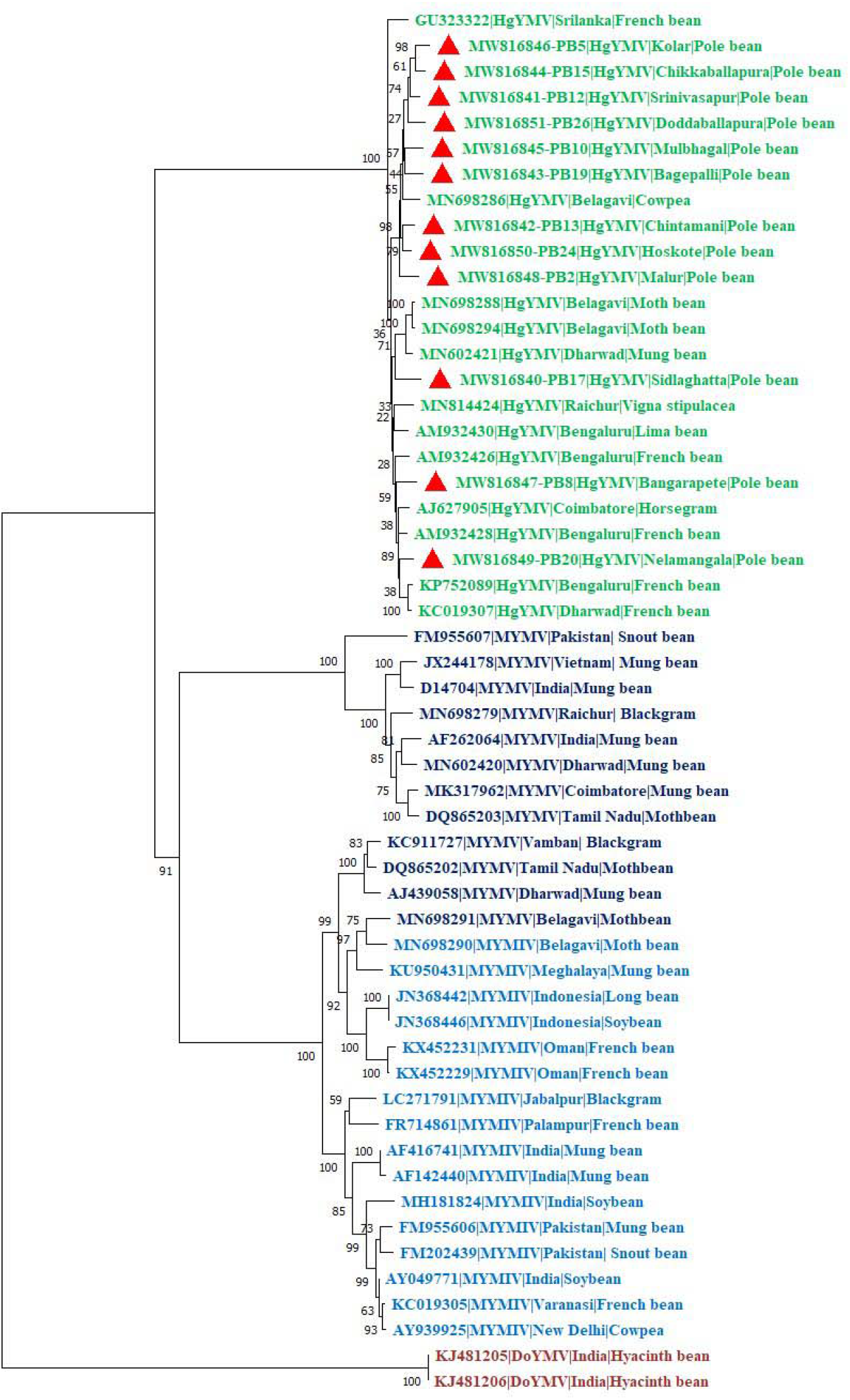
Phylogenetic relationship of nucleotide sequences of DNA-B component of 12 HgYMV isolates from pole bean with other selected 42 begomoviruses infecting legumes bye Neighbour Joining method using MEGA X with 1000 bootstrap replicates

Further, comparison of DNA-B ORFs (BC1 and BV1) of 12 HgYMV isolates from pole bean with other selected 42 begomoviruses infecting legume revealed that they share maximum nt and aa identity with the HgYMV isolates present in the database and low nt and aa identity with other legume infecting begomoviruses (MYMIV, MYMV and DoYMV).

Non-coding region of 12 HgYMV isolates ranged from 240-300 nt in length in DNA-A and DNA-B. Analysis of IR in DNA-A showed 40.10-99.60 per cent identity with HgYMV, HgYMV isolate of horsegram, MN602403 is an outlier and only this isolate has shown low homology of 40.10 per cent with isolates of polebean. and low nt identity with other legume infecting begomoviruses (MYMIV, MYMV, VbGMV, DoYMV, FbLCV, ToLCGV and TbCSV) (Table S5). Whereas, in DNA-B, these 12 HgYMV isolates shared sequence identity of 92.30 to 99.10 per cent with HgYMV isolates and low nt identity with MYMIV, MYMV and DoYMV infecting legume available in the database (Table S6). The two components (DNA-A and DNA B) of bipartite genomes share a CR of approximately 240 nt having an inverted repeat sequence that can potentially form a hairpin-like structure, form a loop having invariant nonanucleotide TAATATTAC and tandemly repeated motifs called as iterons, which are needed for the recognition of origin of replication.

### 3.4 GC-plot and recombination analysis

The GC plot represents the proportion of guanine and cytosine in a stretch of DNA. As the DNA-A and DNA-B of all the 12 HgYMV isolates from pole bean have shown high sequence similarity between themselves, the GC-plot graph of one of the isolates (PB-8) has been represented in the figure 5. The GC plot graph of all the isolates has been depicted in Figure S3. The inner most track with the bars represents the above and below average GC content distribution of the genome. In DNA-A, low GC content was observed in half of the region of the genome encoding AV1 (CP) protein and some regions of the genome encoding AV2, AV3 and AC1 proteins. Some parts of the genome encoding AV1 (CP) and complete regions of the genome encoding AC2 and AC4 proteins were shown to have high GC content. In DNA-B, low GC content was observed in some parts of the genome encoding BC1 and BV1 proteins (Fig. 5). However, the less intensity of GC content was more in the BC1 protein encoding region compared to BV1. As some of the literature survey indicates the possibility of high recombination in the low GC region (Brown 2015, Ninh, 2013 Robinson et al., 2013), the recombination analysis was performed.

**Figure 5:**
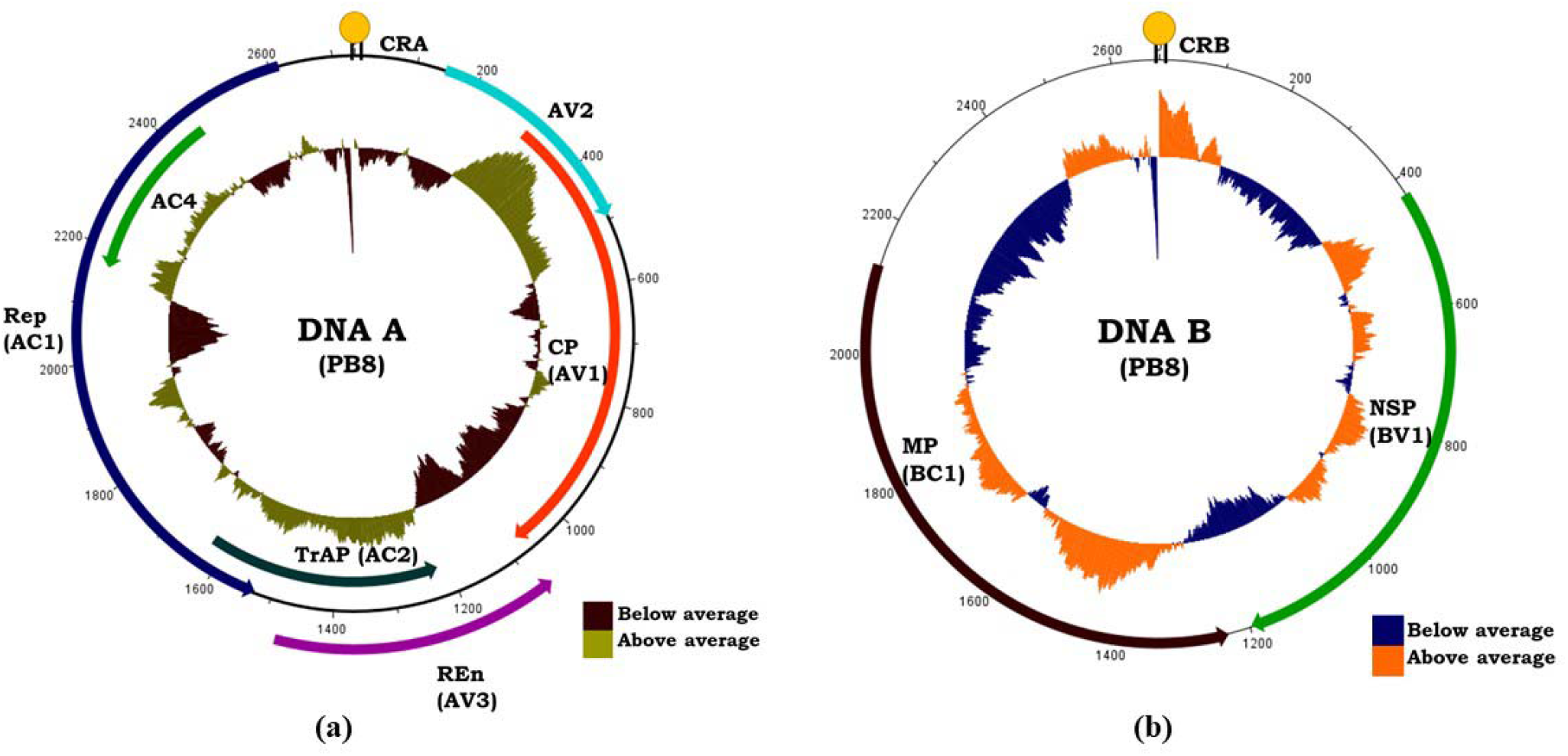
The GC plot graph for (a) DNA-A and (b) DNA-B. The outer most track/circle represents the nucleotide position in the genome. The arrows with different colour code represent the ORFs encoded by DNA-A (a) and DNA-B (b). The inner most circle with coloured bars depicts the above and below average GC content. The graph was generated using Artemis DNA plotter version 18.1.0.

The neighbor-network analysis (Split-tree) of 12 HgYMV isolates from pole bean (DNA-A and DNA-B) displayed reticulate topologies indicating several potential recombination events among putative recombination in the genome (DNA-A and DNA-B) in these isolates (Fig. S4 and S5).

Recombination breakpoint analysis using RDP4 showed intraspecific recombination among the pole bean isolates. Recombination breakpoints detected by more than three methods are considered to be significant recombinants. The analysis of DNA-A of 12 HgYMV isolates revealed that among 12 pole bean HgYMV isolates, only 2 pole bean HgYMV isolates (PB8 and PB13) had shown recombinant breakpoints in more than three methods (Table 1, Fig. 6).

**Figure 6:**
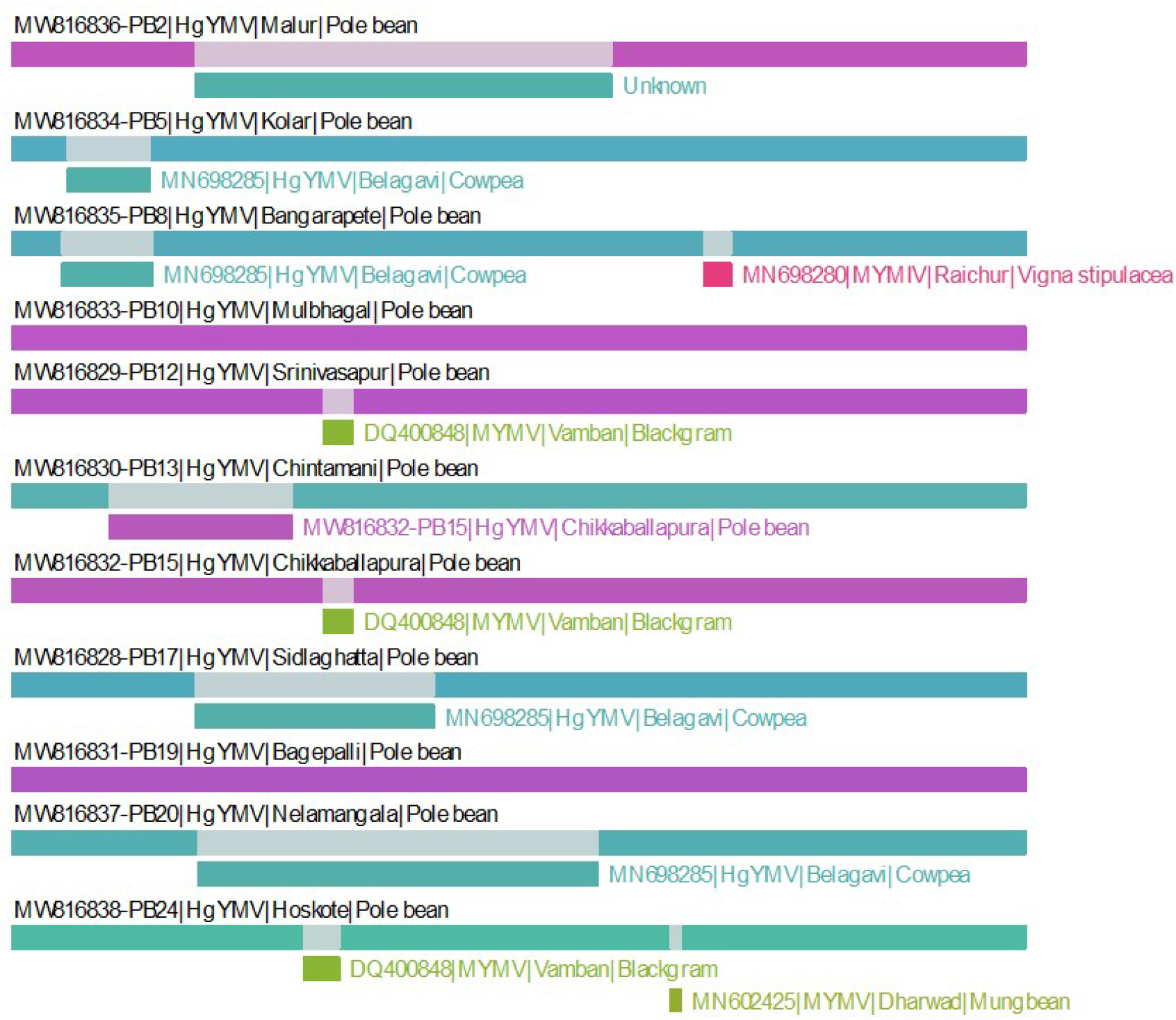
Recombination breakpoint analysis of DNA-A component of 12 HgYMV isolates from pole bean with other selected 50 begomoviruses infecting legumes using RDP, GeneCov, MaxChi, ChimaEra, SiScan and 3Seq integrated in RDP4

**Table 1:** Breakpoint analysis of DNA-A component of 12 HgYMV isolates from pole bean with their putative parent sequences.

| HgYMV isolates | Break point (Beginning-Ending) | Major Parent | Minor Parent | P-Values (0.05) |  |  |  |  |  |
| --- | --- | --- | --- | --- | --- | --- | --- | --- | --- |
|  |  |  |  | RDP | GENECOV | Max Chi | Chimaera | SiScan | 3 Seq |
| MW816836 PB2 | 613-1476 | MW816828-PB17 HgYMV Pole bean | MN698285 HgYMV Cowpea | NS | NS | $3.001 \times 10^{-06}$ | NS | NS | NS |
| MW816834 PB5 | 219-463 | MW816837-PB20 HgYMV Pole bean | MN698285 HgYMV Cowpea | NS | NS | NS | NS | $2.521 \times 10^{-03}$ | NS |
| MW816835 PB8 | 219-463 | MW816837-PB20 HgYMV Polebean | MN698285 HgYMV Cowpea | NS | NS | NS | NS | $2.521 \times 10^{-03}$ | NS |
| | 2584-2641 | MW816834-PB5 HgYMV Pole bean | MN698280 MYMIV Vigna stipulacea | NS | $3.825 \times 10^{-21}$ | $8.095 \times 10^{-05}$ | $3.075 \times 10^{-02}$ | NS | NS |
| MW816833 PB10 | No recombinant |  |  |  |  |  |  |  |  |
| MW816829 PB12 | 1118-1352 | MN698289 MYMIV Moth bean | DQ400848 MYMV Black gram | NS | $5.045 \times 10^{-01}$ | NS | NS | $5.604 \times 10^{-04}$ | NS |
| MW816830 PB13 | 389-887 | MN698285 HgYMV Cow pea | MW816832-PB15 HgYMV Pole bean | $3.329 \times 10^{-05}$ | $1.342 \times 10^{-03}$ | $3.340 \times 10^{-05}$ | $9.910 \times 10^{-05}$ | $2.144 \times 10^{-07}$ | NS |
| MW816832 PB15 | 1125-1218 | MN698289 MYMIV Moth bean | DQ400848 MYMV Black gram | NS | $5.045 \times 10^{-01}$ | NS | NS | $5.604 \times 10^{-04}$ | NS |
| MW816828 PB17 | 578-1570 | MN698285 HgYMV Cow pea | MW816829-PB12 HgYMV Pole bean | NS | NS | $8.805 \times 10^{-06}$ | $4.572 \times 10^{-06}$ | NS | NS |
| MW816831 PB19 | No recombinant |  |  |  |  |  |  |  |  |
| MW816837 PB20 | 613-1921 | MW816829-PB12 HgYMV Pole bean | MN698285 HgYMV Cow pea | NS | NS | $1.071 \times 10^{-03}$ | NS | $5.267 \times 10^{-07}$ | NS |
| MW816838 PB24 | 1072-1218 | MN698289 MYMIV Moth bean | DQ400848 MYMV Black gram | NS | $5.045 \times 10^{-01}$ | NS | NS | $5.604 \times 10^{-04}$ | NS |
| | 2473-2501 | MN698293 MYMV Moth bean | MN602425 MYMV Mung bean | NS | $2.822 \times 10^{-06}$ | NS | NS | NS | NS |
| MW816839 PB26 | 1072-1218 | MN698289 MYMIV Moth bean | DQ400848 MYMV Black gram | NS | $5.045 \times 10^{-01}$ | NS | NS | $5.604 \times 10^{-04}$ | NS |
| | 1280-1353 | AM932427 HgYMV Cow pea | KC911721 MYMV Black gram | $9.658 \times 10^{-14}$ | $9.659 \times 10^{-13}$ | NS | NS | NS | NS |

Further, analysis also revealed that, most part of DNA-A fragments (AC1) of MW816835-PB8 isolate might have descended from recombination between MW816834-PB5 (HgYMV) and MN698280 (MYMIV) infecting pole bean and *Vigna stipulacea*, as major and minor parents, respectively. Whereas in case of MW816830-PB 13 isolate, most part of DNA-A fragments (AV1 and AV2) might have descended from the recombination between MN698289 and MW816832-PB15 (HgYMV) infecting cowpea and pole bean as major and minor parents, respectively (Table 1, Fig. 6).

A similar analysis was carried out for DNA-B associated with the 12 HgYMV isolates from pole bean. Analysis revealed the presence of intraspecific recombination in all 12 pole bean isolates). In case of PB2 isolate, two recombination breakpoints were detected with most part (BV1, BC1) of DNA-B component derived from the recombination between HgYMV (KP752089) and HgYMV (MW816844 -PB15) infecting french bean and pole bean, respectively as major and minor parents. While the second breakpoint, most part (BC1) of DNA-B component might have originated from the recombination between HgYMV isolates (MN698294, MW816845-PB10) infecting moth bean and pole bean, respectively as major and minor parents. In the case of PB 5 three recombination events were detected. In PB5 (first break), PB12, PB15 and PB17 isolates with most part (BV1, IR) of DNA-B component derived from the recombination between HgYMV (AJ627905 and GU323322) infecting horsegram and french bean as major and minor parents. The PB 17 isolate showed two recombination events. While the second breakpoint in both PB5 and PB17 isolates might have descended from recombination between AM932428 (HgYMV) and MW816844-PB15 (HgYMV) infecting french bean and pole bean and whereas in the third breakpoint of PB5 and PB10 isolates, most part (BV1, BC1) of DNA-B component derived from MW816844-PB15 and MW816843-PB19 isolates infecting pole bean as major and minor parents, respectively. In PB19, (BV1 and IR) DNA-B descended from HgYMV (MW816840-PB17 and MW816844-PB15) infecting pole bean as major and minor parents, respectively (Table 2, Fig 7). The overall recombination analysis showed that, DNA-A form two isolates (PB8 and PB13) and DNA-B associated with PB2, PB5, PB10, PB12, PB13, PB15, PB17 and PB19 isolates are considered to be recombinants.

**Figure 7:**
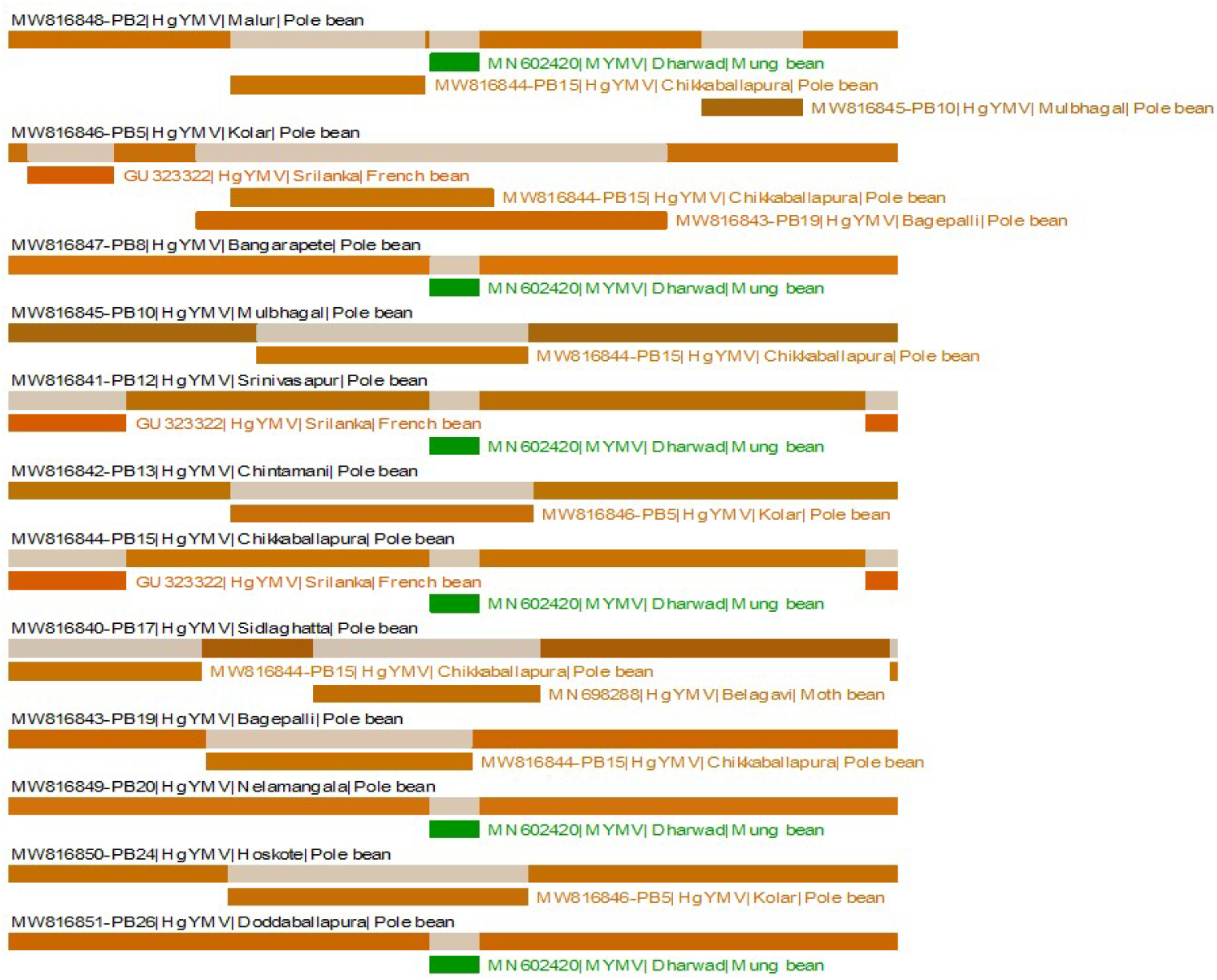
Recombination breakpoint analysis of DNA-B component of 12 HgYMV isolates from pole bean with other selected 42 begomoviruses infecting legumes using RDP, GeneCov, MaxChi, Chimaera, SiScan and 3Seq integrated in RDP4

**Table 2:** Breakpoint analysis of DNA-B component of 12 HgYMV isolates from pole bean with their putative parent sequences.

| HgYMV isolates | Break point (Beginning-Ending) | Major Parent | Minor Parent | P-Values (0.05) |  |  |  |  |  |
| --- | --- | --- | --- | --- | --- | --- | --- | --- | --- |
|  |  |  |  | RDP | GENECOV | Max Chi | Chimaera | SiScan | 3 Seq |
| MW816848 PB2 | 792-1334 | KP752089 HgYMV French bean | MW816844- PB15 HgYMV Pole bean | $3.713 \times 10^{-09}$ | $9.989 \times 10^{-08}$ | $1.745 \times 10^{-07}$ | $1.746 \times 10^{-08}$ | $2.965 \times 10^{-09}$ | NS |
| | 1307-1460 | AJ439058 MYMV Mung bean | MN602420 MYMV Mung bean | NS | NS | $9.633 \times 10^{-03}$ | $2.771 \times 10^{-02}$ | NS | NS |
| | 1909-2282 | MN698294 HgYMV Moth bean | MW816845- PB10 HgYMV Pole bean | $2.576 \times 10^{-03}$ | $2.096 \times 10^{-04}$ | $8.397 \times 10^{-04}$ | $2.494 \times 10^{-03}$ | $1.128 \times 10^{-05}$ | NS |
| MW816846 PB5 | 128-412 | AJ627905 HgYMV Horse gram | GU323322 HgYMV French bean | $2.458 \times 10^{-03}$ | $2.516 \times 10^{-03}$ | NS | NS | $5.371 \times 10^{-04}$ | NS |
| | 562-1387 | AM932428 HgYMV French bean | MW816844- PB15 HgYMV Pole bean | $4.534 \times 10^{-02}$ | $2.097 \times 10^{-05}$ | $6.969 \times 10^{-05}$ | $4.955 \times 10^{-05}$ | $3.294 \times 10^{-11}$ | NS |
| | 563-2165 | MW816844- PB15 HgYMV Pole bean | MW816843- PB19 HgYMV Pole bean | $1.677 \times 10^{-03}$ | $1.721 \times 10^{-18}$ | $4.298 \times 10^{-05}$ | $1.154 \times 10^{-02}$ | NS | NS |
| MW816847 PB8 | 1307-1460 | AJ439058 MYMV Mung bean | MN602420 MYMV Mung bean | NS | NS | $9.633 \times 10^{-03}$ | $2.771 \times 10^{-02}$ | NS | NS |
| MW816845 PB10 | 563-2165 | MW816844- PB15 HgYMV Pole bean | MW816843- PB19 HgYMV Pole bean | $1.677 \times 10^{-03}$ | $1.721 \times 10^{-18}$ | $4.298 \times 10^{-05}$ | $1.154 \times 10^{-02}$ | NS | NS |
| MW816841 PB12 | 2517-412 | AJ627905 HgYMV Horse gram | GU323322 HgYMV French bean | $2.458 \times 10^{-03}$ | $2.516 \times 10^{-03}$ | NS | NS | $5.371 \times 10^{-04}$ | NS |
| | 1307-1460 | AJ439058 MYMV Mung bean | MN602420 MYMV Mung bean | NS | NS | $9.633 \times 10^{-03}$ | $2.771 \times 10^{-02}$ | NS | NS |
| MW816842 PB13 | 768-1706 | AM932428 HgYMV French bean | MW816846- PB5 HgYMV Pole bean | $8.523 \times 10^{-11}$ | $5.275 \times 10^{-09}$ | $7.157 \times 10^{-11}$ | $1.830 \times 10^{-10}$ | $7.968 \times 10^{-15}$ | NS |
| MW816844 PB15 | 2517-412 | AJ627905 HgYMV Horse gram | GU323322 HgYMV French bean | $2.458 \times 10^{-03}$ | $2.516 \times 10^{-03}$ | NS | NS | $5.371 \times 10^{-04}$ | NS |
| | 1307-1460 | AJ439058 MYMV Mung bean | MN602420 MYMV Mung bean | NS | NS | $9.633 \times 10^{-03}$ | $2.771 \times 10^{-02}$ | NS | NS |
| MW816840 PB17 | 81-494 | AJ627905 HgYMV Horse gram | MW816844- PB15 HgYMV Pole bean | $1.111 \times 10^{-07}$ | $1.692 \times 10^{-07}$ | $4.854 \times 10^{-05}$ | $4.833 \times 10^{-03}$ | $4.626 \times 10^{-05}$ | NS |
| | 753-1772 | AM932428 HgYMV French bean | MN698288 HgYMV Moth bean | NS | NS | $1.295 \times 10^{-03}$ | $1.327 \times 10^{-04}$ | $6.149 \times 10^{-07}$ | NS |
| MW816843 PB19 | 577-1661 | MW816840- PB17 HgYMV Pole bean | MW816844- PB15 HgYMV Pole bean | $1.085 \times 10^{-06}$ | $2.613 \times 10^{-06}$ | $9.049 \times 10^{-11}$ | $3.877 \times 10^{-06}$ | $3.423 \times 10^{-15}$ | NS |
| MW816849 PB20 | 1307-1460 | AJ439058 MYMV Mung bean | MN602420 MYMV Mung bean | NS | NS | $9.633 \times 10^{-03}$ | $2.771 \times 10^{-02}$ | NS | NS |
| MW816850 PB24 | 1307-1460 | AJ439058 MYMV Mung bean | MN602420 MYMV Mung bean | NS | NS | $9.633 \times 10^{-03}$ | $2.771 \times 10^{-02}$ | NS | NS |
| MW816851 PB26 | 1307-1460 | AJ439058 MYMV Mung bean | MN602420 MYMV Mung bean | NS | NS | $9.633 \times 10^{-03}$ | $2.771 \times 10^{-02}$ | NS | NS |

## 4 Discussion

Begomoviruses infecting legumes are unique and genetically different from other begomoviruses infecting the rest of agricultural and horticultural crops (Agnihotri et al., 2019). These DNA viruses are endemic in distribution and present only in Indian subcontinent and Southeast Asia and proposed as a distinct group ‘‘Legumovirus’’ known to infect grain legumes and weeds. DNA viruses associated with yellow mosaic disease (YMD) of legumes are genetically isolated, have a narrow host range and do not have opportunities to undergo genetic re-assortment or recombination, which occur frequently in other begomoviruses having wider host range. Despite this barrier, they continue to cause disease in epidemic proportion. In the present study, a roving survey was carried out to know the prevalence of YMD of pole bean, which is extensively cultivated in different parts of Karnataka, India. In all the surveyed regions, ubiquitous prevalence of YMD in pole bean was observed. In most of the fields, infected plants exhibited symptoms of vein clearing, mosaic, downward curling, yellowing, stunted growth and deformed fruits with immature pods. The PDI of YMD ranged from 6.02 to 80.74 per cent in different surveyed areas. The available literature on the survey of YMD also reveals its ubiquitous presence all over the country (Nariani, 1960; Nene, 1978). Further, Archith et al. (2017) carried out similar survey work on YMD of french bean and recorded disease incidence ranging from 5.88 to 28.54 per cent. The reason behind this variation in the occurrence of YMD in different places may be due to weather parameters like temperature, relative humidity, vector population, feeding behavior and infected surrounding crops that help in the transmission of the virus and also resistant cultivars.

HgYMV was first recorded on horsegram by William et al. (1968). Subsequently it was reported on many crops that includes common bean (Monger et al., 2010), horsegram (Muniyappa et al., 2008), lima bean (Abarshi et al., 2017), soybean and *Hedyotis corymbosa* (Rienzie et al., 2016). In the current study, the pole bean leaf samples collected from different locations of Karnataka, India, were confirmed for the presence of begomovirus by PCR detection. Complete genome sequence analysis showed that the begomovirus associated with YMD of pole bean as HgYMV, which shared more than 91 per cent nt identity with several HgYMV isolates infecting different leguminous crops *viz*., cowpea, french bean, moth bean, lima bean, horsegram and low sequence identity (< 85%) with several isolates of MYMIV, MYMV, VbGMV, DoYMV FbLCV, ToLCGV and TbCSV infecting different legumes in India.

Analysis of the CR sequences of DNA-A and DNA-B sequences of the 12 HgYMV isolates from pole bean revealed that they possessed more homology with several isolates of HgYMV infecting cowpea, french bean, moth bean, lima bean and horsegram. The IR contains a predicted stem-loop sequence with a conserved nonanucleotide sequence (TAATATTAC) in the loop, which acts as the origin of viral DNA replication and can be found in the majority of geminiviruses characterized so for (Heyraud et al., 1993)

The iteron sequences of DNA-A and DNA-B sequences identified in the stem loop regions of 12 HgYMV isolates from pole bean were identical with several isolates of HgYMV infecting cowpea, french bean, moth bean, lima bean, horsegram. The iteron sequence is recognized as the site for sequence-specific binding Rep protein which results in the initiation of viral replication (Heyraud et al., 1993) Any modifications occurring in this site with respect to sequence, size or number of iterons, or even a point mutation depending on the viral species, adversely affects the binding ability of Rep protein both *in vivo* and *in vitro* (Chatterji et al., 2000). Literature survey also showed similar ORFs in DNA-A A and DNA-B component of the begomoviruses identified reported so far across the world (Venkataravanappa et al., 2018)

The nt composition of a genome is an important qualitative aspect of genomic architecture and it is widely expressed in the proportion of guanine (G) and cytosine (C) content. To observe the variation in the GC content of HgYMV isolates from pole bean, the GC plot graphs were generated. From the analysis, low GC content was observed in the regions of genome encoding AV1, AV2 and AV3 proteins in DNA-A. Similarly, different intensity of low GC content was observed in the regions of DNA-B encoding BC1 and BV1 proteins. Similar study was carried out by Yogindran et al. (2021), wherein, the low GC content was observed in the genomic region encoding AV1 protein. In human adenovirus, the emergence of highly virulent strains was found to be due to the homologous recombination in the low GC rich region (Robinson et al., 2013). In order to find such a correlation between GC content and recombination, the recombination analysis was performed.

Evolution in plant viruses may occur through various factors that include virus-vector and virus-host interactions, mixed infections and also through the high level of viral replication (Padidam et al., 1995). In begomoviruses, most frequent and major diversification mechanism occurs through recombination (Padidam et al., 1995: Lopez et al., 2009) and resulting in emergence of novel viruses. However, mixed infection, adaptability and extension of its host range are the prerequisites for recombination. Recombination is a common phenomenon that occurs at species level (Lozano et al., 2009) or at strainal level (Fondong et al., 2000). Recombination analysis showed that 12 isolates associated with YMD disease of pole bean are a recombinant variant of previously reported viruses. Recombination events observed in both DNA-A and DNA-B genomic regions having low GC rich regions. Stability of the GC rich region in an organism is due to triple hydrogen bonding between the nucleotides and may be this could be one of the reasons for lesser recombination (Brown, 2015; Ninh, 2013; Robinson et al., 2013). Recombination is a rapid process to create new genomes with adaptive advantages, which could accelerate their evolution, favouring expansion of the host range and therefore the emergence of novel diseases (Venkataravanappa et al., 2014).

## 6 Contributions

BJ, HDV, MN, MM performed the biological experiment, SH, VV and KSS involved in the bioinformatics analysis CRJB, D and CNL involved in experimental design, CNL conceptualized work and provided the overall direction, all authors are involved in literature mining and manuscript preparation. All authors read and approved the final manuscript.

## 7 Ethics declarations

### 7.2 Research involving human participants and/or animals

This article does not contain any studies with human participants or animals performed by any of the authors.

### 7.3 Informed consent

Informed consent was obtained from all individual participants included in the study.

## 10 Supplementary Figure Legends

**Figure S1:** Graphical representation of percentage pair wise genome scores and nucleotide identity plot of DNAAA component of 12 HgYMV isolates from pole bean with selected sequences of 50 begomovirus isolates prepared using Sequence Demarcation Tool version 1.2 (SDTv1.0)

**Figure S2:** Graphical representation of percentage pair wise genome scores and nucleotide identity plot of DNA-B component of 12 HgYMV isolates from pole bean, with other 50 begomovirus isolates sequences-prepared using Sequence Demarcation Tool version 1.2 (SDTv1.2)

**Figure S3:** The GC plot graph of all the 12 HgYMV isolates infecting pole beans (a) DNA-A and (b) DNA-B. The outermost circle represents the nucleotide position of the circular genome. The inner circle with colour coded bars depict the above and below average GC content of the respective genome. The graphs were generated with the aid of Artemis DNA plotter version 18.1.0.

**Figure S4:** A neighbour-net generated for sequences of DNA-A component of 12 HgYMV isolates from pole bean with other selected 50 begomoviruses infecting legumes.

**Figure S5:** A neighbour-net generated for sequences of DNA-B component of 12 HgYMV isolates from pole bean with other selected 42 begomoviruses infecting legumes.

## Supplementary Table Legends

**Table S1:** List of selected begomoviruses used in the study for comparison analysis of DNA-A component of the 12 pole bean HgYMV isolates

**Table S2:** List of selected begomoviruses used in the study for comparison analysis of DNA-B component of the 12 pole bean HgYMV s isolates

**Table S3:** Percent disease incidence of yellow mosaic disease of pole bean in Karnataka during 2019-2020

**Table-S4:** Nucleotide and amino acid identity of DNA-A of 12 HgYMV isolates from pole bean with other selected 50 begomoviruses infecting legumes

**Table S5:** Nucleotide and amino acid identity of DNA-B of 12 HgYMV isolates from pole bean with other selected 42 begomoviruses infecting legumes

## References

Abarshi, M. M., Abubakar, A. L., Garba, A., Mada, S. B., Ibrahim, A. B., & Maruthi, M. N. (2017). Molecular detection and characterization of *Horsegram yellow mosaic virus* (HgYMV) infecting lima bean (*Phaseolus lunatus*) in India. Nigerian Journal of Biotechnology, 33, 41–48.

Agnihotri, A. K., Mishra, S. P., Ansar, M., Tripathi, R. C., Singh, R., & Akram, M. (2019). Molecular characterization of *Mungbean yellow mosaic* India virus infecting tomato (*Solanum lycopersicum* L.). Australasian Plant Pathology, 48(2), 159–165.

Ansar, M., Agnihotri, A. K., Akram, M., & Bhagat, A. P. (2019). First report of *Tomato leaf curl Joydebpur virus* infecting French bean (*Phaseolus vulgaris* L.). Journal of General Plant Pathology, 85(6), 444–448. doi:10.1007/s10327-019-00867-5

Archith, T.C., Devappa, V., Prashant & Naganur Priya, I., (2017). Status of *Mungbean Yellow Mosaic Virus* (MYMV) on French bean in different agro-climatic zones of Karnataka, India. International Journal of Agriculture Sciences, ISSN: 0975-3710 & E-ISSN: 0975-9107, Volume 9, Issue 11, pp.-4015–4019

Brown JK, Zerbini FM, Navas-Castillo J, Moriones E, Ramos-Sobrinho R, Silva JC, Fiallo-Olivé E, Briddon RW, Hernández-Zepeda C, Idris A, Malathi VG, Martin DP, Rivera-Bustamante R, Ueda S, Varsani A. (2015). Revision of Begomovirus taxonomy based on pairwise sequence comparisons. Archives of Virology. 160(6):1593–619. doi: 10.1007/s00705-015-2398-y.

Chatterji, A., Chatterji, U., Beachy, R. N., & Fauquet, C. M. (2000). Sequence parameters that determine specificity of binding of the replication-associated protein to its cognate site in two strains of *Tomato leaf curl virus New Delhi*. Virology, 273(2), 341–350.

De Barro, P. J., Liu, S. S., Boykin, L. M., & Dinsdale, A. B. (2011). *Bemisia tabaci*: a statement of species status. Annual Review of Entomology, 56, 1–19. doi: 10.1146/annurev-ento-112408-085504.

Doyle, J.J. and J.L. Doyle, 1990. Isolation of plant DNA from fresh tissue. Focus, 12: 13–15.

Euesden, J., Lewis, C. M., & O’Reilly, P. F. (2015). PRSice: polygenic risk score software. Bioinformatics, 31(9), 1466–1468. doi: 10.1093/bioinformatics/btu848.

Fauquet, C. M., Bisaro, D. M., Briddon, R. W., Brown, J. K., Harrison, B. D., Rybicki, E. P., & Stanley, J. (2003). Revision of taxonomic criteria for species demarcation in the family *Geminiviridae*, and an updated list of begomovirus species. Archives of Virology, 148(2), 405–420. doi: 10.1007/s00705-002-0957-5.

Fondong, V. N., Pita, J. S., Rey, M. E. C., De Kochko, A., Beachy, R. N., & Fauquet, C. M. (2000). Evidence of synergism between *African cassava mosaic virus* and a new double-recombinant geminivirus infecting cassava in Cameroon. Journal of General Virology, 81(1), 287–297.

Gallo, C., Renzi, P., Loizzo, S., Loizzo, A., Piacente, S., Festa, M., … & Capasso, A. (2010). Potential therapeutic effects of vitamin E and C on placental oxidative stress induced by nicotine: an in vitro evidence. The Open Biochemistry Journal, 4, 77. doi: 10.2174/1874091X01004010077.

Gámez-Jiménez, C., Romero-Romero, J. L., Santos-Cervantes, M. E., Leyva-Lopez, N. E., & Mendez-Lozano, J. (2009). Tomato as a natural new host for *Tomato yellow leaf curl virus* in Sinaloa, Mexico. Plant disease, 93(5), 545–545.

Hanley-Bowdoin, L., Bejarano, E. R., Robertson, D., & Mansoor, S. (2013). Geminiviruses: masters at redirecting and reprogramming plant processes. Nature Reviews Microbiology, 11(11), 777–788. doi: 10.1038/nrmicro3117.

Hanley-Bowdoin, L., Settlage, S. B., Orozco, B. M., Nagar, S., & Robertson, D. (1999). Geminiviruses: models for plant DNA replication, transcription, and cell cycle regulation. Critical Reviews in Plant Sciences, 18(1), 71–106.

Heyraud, F., Matzeit, V., Kammann, M., Schaefer, S., Schell, J., & Gronenborn, B. (1993). Identification of the initiation sequence for viralustrand DNA synthesis of *Wheat dwarf virus*. The EMBO Journal, 12(11), 4445–4452.

Huson, D. H., & Bryant, D. (2006). Application of phylogenetic networks in evolutionary studies. Molecular biology and evolution, 23(2), 254–267. doi: 10.1093/molbev/msj030.

Ilyas, M., Qazi, J., Mansoor, S., & Briddon, R. W. (2009). Molecular characterization and infectivity of a “Legumovirus” (genus Begomovirus: family *Geminiviridae*) infecting the leguminous weed *Rhynchosia minima* in Pakistan. Virus Research, 145(2), 279–284. doi: 10.1016/j.virusres.2009.07.018.

Jeevan, B., Nagaraju, N., & Basavaraj, S. (2015). Molecular detection and partial characterization of begomovirus associated yellow mosaic virus disease of pole bean (*Phaseolus vulgaris* L.) in Southern India. Journal of Pure and Applied Microbiology. 9(1), p. 227–235

Kamaal, N., Akram, M., & Agnihotri, A. K. (2015). Molecular Evidence for the Association of *Tomato leaf curl Gujarat virus* with a Leaf Curl Disease of *Phaseolus vulgaris* L. Journal of Phytopathology, 163(1), 58–62. doi: 10.1007/s10327-019-00867-5

Kamaal, N., Akram, M., Pratap, A., & Yadav, P. (2013). Characterization of a new begomovirus and a beta satellite associated with the leaf curl disease of French bean in northern India. Virus Genes, 46(1), 120–127. doi: 10.1007/s11262-012-0832-8

Kumar, S., Stecher, G., Li, M., Knyaz, C., & Tamura, K. (2018). MEGA X: molecular evolutionary genetics analysis across computing platforms. Molecular biology and evolution, 35(6), 1547.

Martin, D. P., Murrell, B., Golden, M., Khoosal, A., & Muhire, B. (2015). RDP4: Detection and analysis of recombination patterns in virus genomes. Virus Evolution, 1(1). doi: 10.1093/ve/vev003

Monger, W. A., Harju, V., Nixon, T., Bennett, S., Reeder, R., Kelly, P., & Ariyarathne, H. M. (2010). First report of *Horsegram yellow mosaic virus* infecting *Phaseolus vulgaris* in Sri Lanka. New Disease Reports, 21(16), 2044–0588.

Muhire, B. M., Varsani, A., & Martin, D. P. (2014). SDT: a virus classification tool based on pairwise sequence alignment and identity calculation. PLOS one, 9(9), e108277. doi: 10.1371/journal.pone.0108277

Muniyappa V, Rajeshwari, R., Bharathan N, Reddy D. N. R., Nolt, B. L., 2008. Isolation and characterization of a geminivirus causing yellow mosaic disease of horsegram (*Macrotyloma uniflorum* [Lam.] Verdc.) in India. Journal of Phytopathology 119, 81–87. [doi:10.1111/j.1439-0434.1987.tb04386.x]

Nariani, T. K. (1960). Yellow mosaic of mung (*Phaseolus aureus* L.). Indian Phytopathol, 13, 24–29.

Nene, Y. L. 1978. A world list of pigeonpea [Cajanus cajan (L.) Millsp.] and chickpea (Cicer arietinum L.) pathogens. ICRISAT Pulse Pathology Progress Report 3. Hyderabad, India.

Nene, Y. L., Hanounik, S. B., Qureshi, S. H., & Sen, B. (1988). Fungal and bacterial foliar diseases of pea, lentil, faba bean and chickpea. In World crops: Cool season food legumes (pp. 577–589). Springer, Dordrecht.

Ninh, A. (2013). Correlation between GC-content and palindromes in randomly generated sequences and viral genomes. arXiv preprint arXiv:1302.5869.

Padidam, M., Beachy, R. N., & Fauquet, C. M. (1995). *Tomato leaf curl geminivirus* from India has a bipartite genome and coat protein is not essential for infectivity. Journal of General Virology, 76(1), 25–35.

Qazi, J., Ilyas, M., Mansoor, S., & Briddon, R. W. (2007). Legume yellow mosaic viruses: genetically isolated begomoviruses. Molecular Plant Pathology, 8(4), 343–348. doi: 10.1111/j.1364-3703.2007.00402.x.

Robinson, C. M., Singh, G., Lee, J. Y., Dehghan, S., Rajaiya, J., Liu, E. B., & Chodosh, J. (2013). Molecular evolution of human adenoviruses. Scientific Reports, 3(1), 1–7. doi: 10.1038/srep01812.

Venkataravanappa, V., Lakshminarayana Reddy, C. N., Jalali, S., & Krishna Reddy, M. (2012). Molecular characterization of distinct bipartite begomovirus infecting bhendi (*Abelmoschus esculentus* L.) in India. Virus Genes, 44(3), 522–535. doi: 10.1007/s11262-012-0732-y.

Venkataravanappa, V., Lakshminarayana Reddy, C. N., Swarnalatha, P., Mahesha, B., Rai, A. B., & Krishna Reddy, M. (2014). Association of *Tomato leaf curl Joydebpur virus* and a betasatellite with leaf curl disease of eggplant. Phytoparasitica, 42(1), 109–120.

Venkataravanappa, V., Reddy, C. L., Saha, S., & Reddy, M. K. (2018). Recombinant *Tomato leaf curl New Delhi virus* is associated with yellow vein mosaic disease of okra in India. Physiological and Molecular Plant Pathology, 104, 108–118.

Venkataravanappa, V., Lakshminarayana Reddy, C. N., Jalali, S., & Krishna Reddy, M. (2012). Molecular characterization of distinct bipartite begomovirus infecting bhendi (Abelmoschus esculentus L.) in India. Virus Genes, 44(3), 522–535.

Williams, F. J., Grewal, J. S., & Amin, K. S. (1968). Serious and new diseases of pulse crops in India in 1966. Plant Disease Reporter, 52(4), 300–304.

Yogindran, S., Kumar, M., Sahoo, L., Sanatombi, K., & Chakraborty, S. (2021). Occurrence of *Cotton leaf curl Multan virus* and associated betasatellites with leaf curl disease of Bhut-Jolokia chillies (*Capsicum chinense* Jacq.) in India. Molecular Biology Reports, 48(3), 2143–2152. doi: 10.1007/s11033-021-06223-1.

Zerbini, F. M., Briddon, R. W., Idris, A., Martin, D. P., Moriones, E., Navas-Castillo, J., & Consortium, I. R. (2017). ICTV virus taxonomy profile: *Geminiviridae*. The Journal of General Virology, 98(2), 131. doi: 10.1099/jgv.0.000738.

Martín-Hernández, I., & Pagán, I. (2022). Gene Overlapping as a Modulator of Begomovirus Evolution. Microorganisms, 10(2), 366.

Robinson, C.M., Singh, G., Lee, J.Y., Dehghan, S., Rajaiya, J., Liu, E.B., Yousuf, M.A., Betensky, R.A., Jones, M.S., Dyer, D.W. and Seto, D. (2013). Molecular evolution of human adenoviruses. Scientific reports, 3(1), pp.1–7.

