## Supplementary tables for "Genetic diversity and evidence of recombination of *Horsegram yellow mosaic virus* infecting pole bean (*Phaseolus vulgaris* L.) from South India"

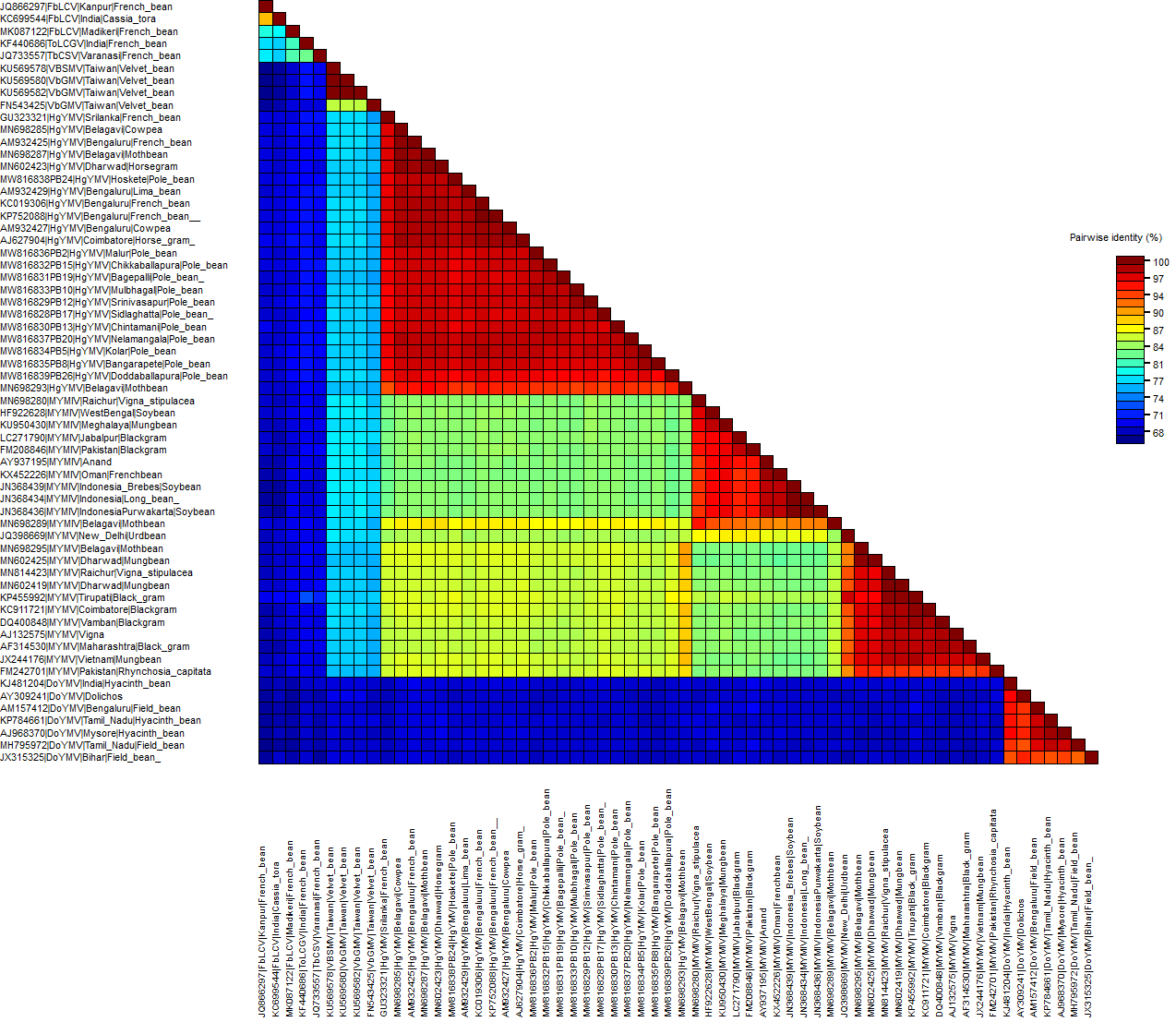

**Figure S1:** Graphical representation of percentage pair wise genome scores and nucleotide identity plot of DNAAA component of 12 HgYMV isolates from pole bean with selected sequences of 50 begomovirus isolates prepared using Sequence Demarcation Tool version 1.2 (SDTv1.0)

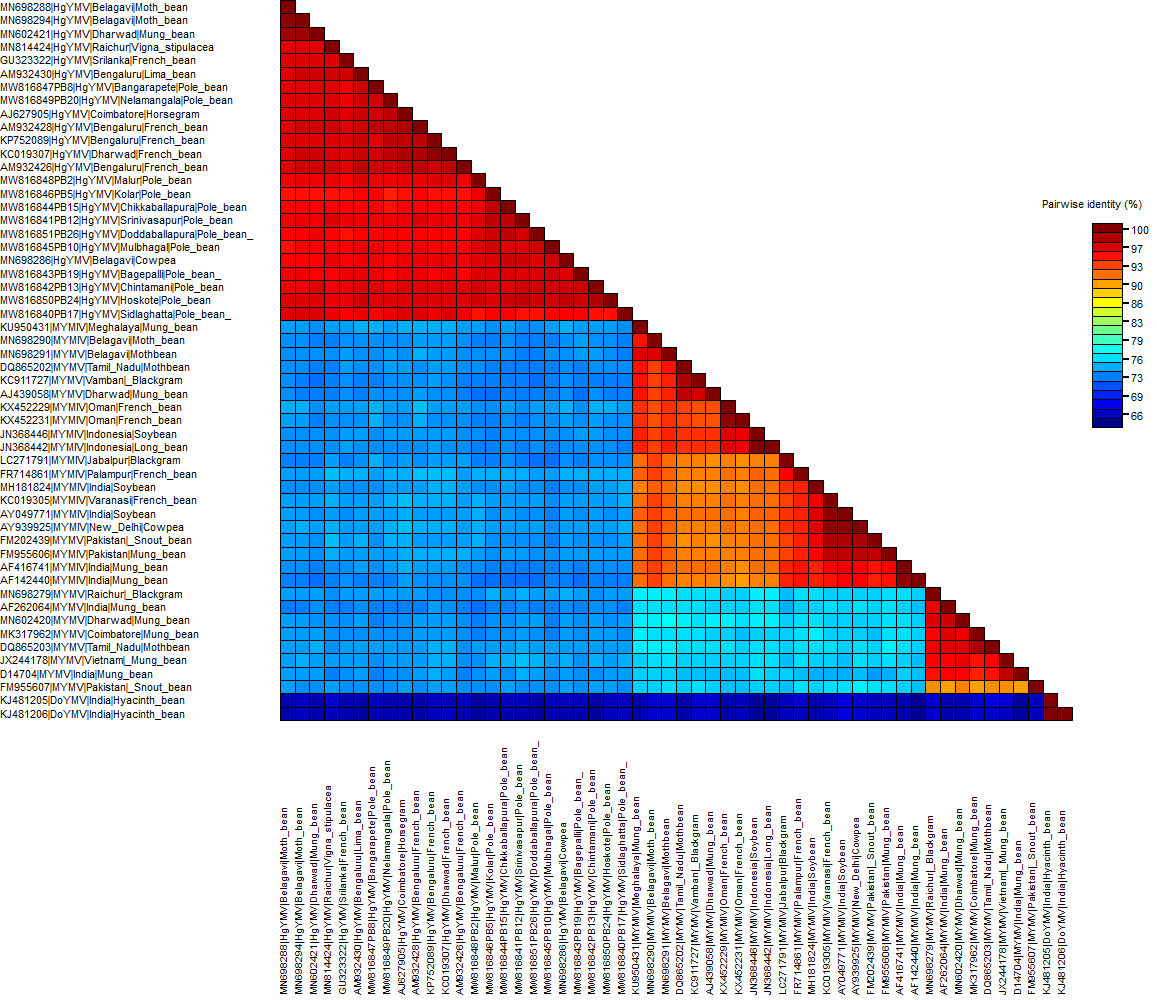

**Figure S2:** Graphical representation of percentage pair wise genome scores and nucleotide identity plot of DNA-B component of 12 HgYMV isolates from pole bean, with other 50 begomovirus isolates sequences- prepared using Sequence Demarcation Tool version 1.2 (SDTv1.2)

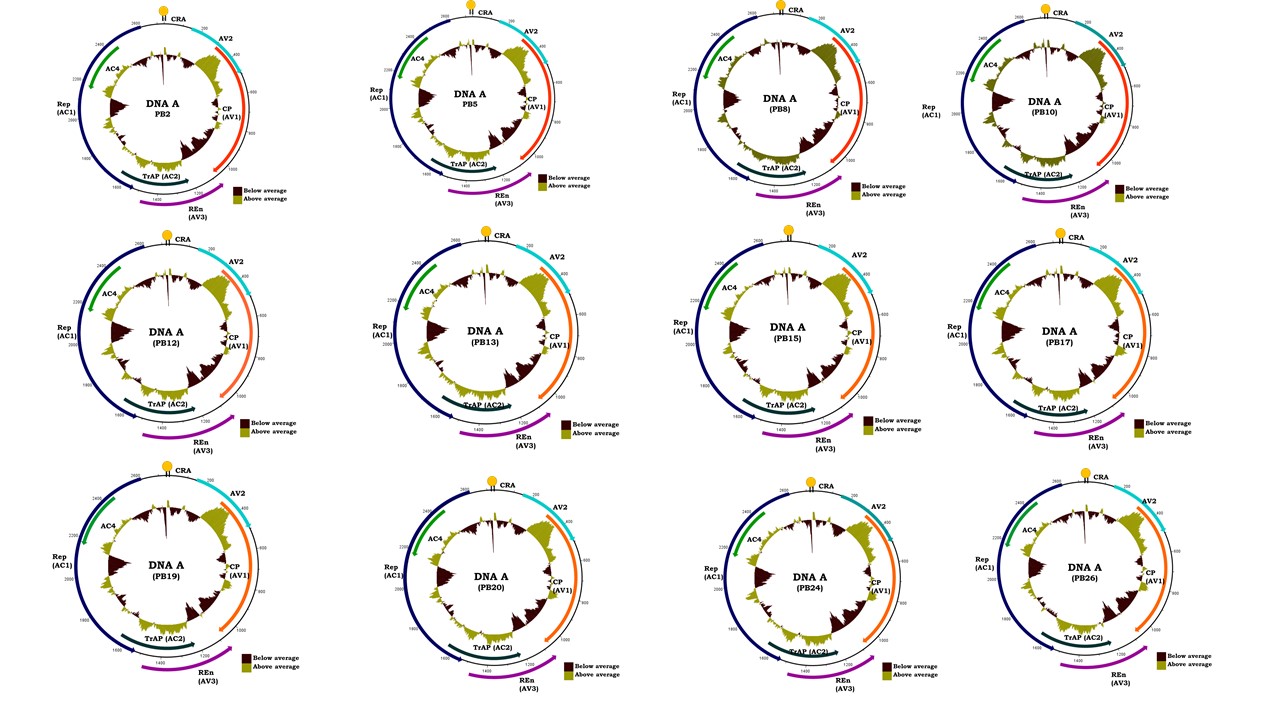

**(a)**

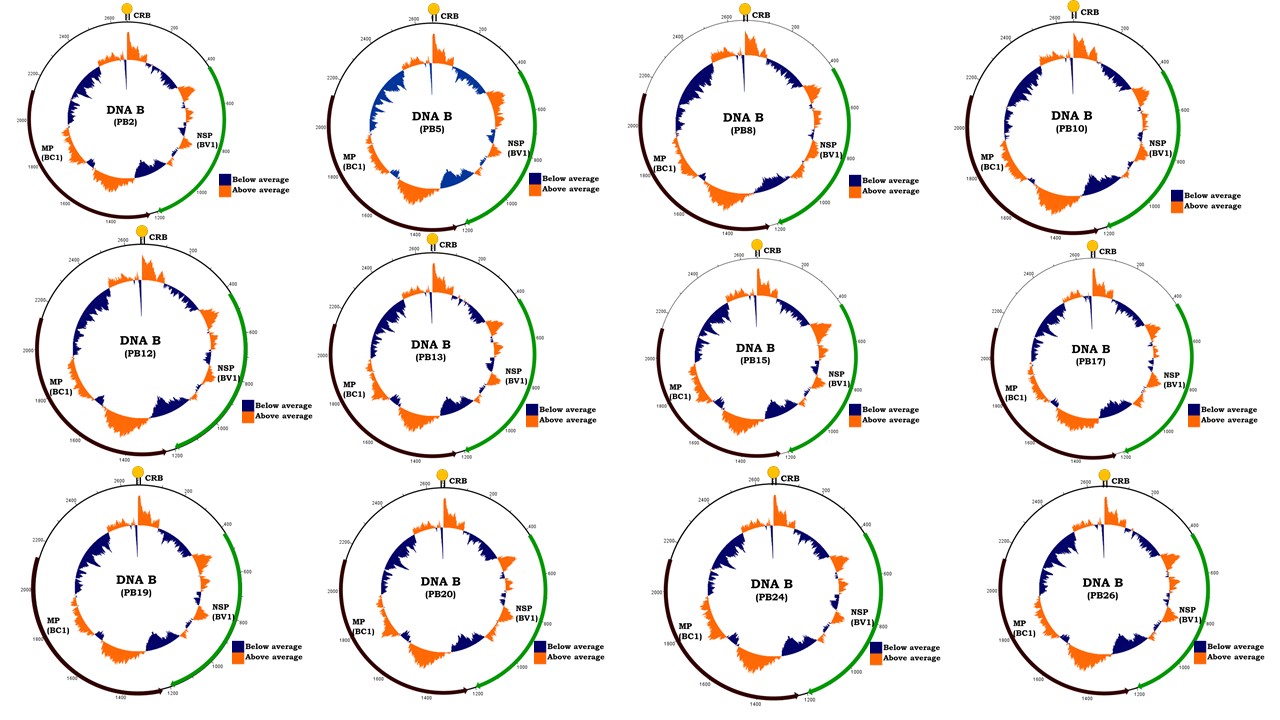
**(b)**

**Figure S3:** The GC plot graph of all the 12 HgYMV isolates infecting pole beans (a) DNA-A and (b) DNA-B. The outermost circle represents the nucleotide position of the circular genome. The inner circle with colour coded bars depict the above and below average GC content of the respective genome. The graphs were generated with the aid of Artemis DNA plotter version 18.1.0.

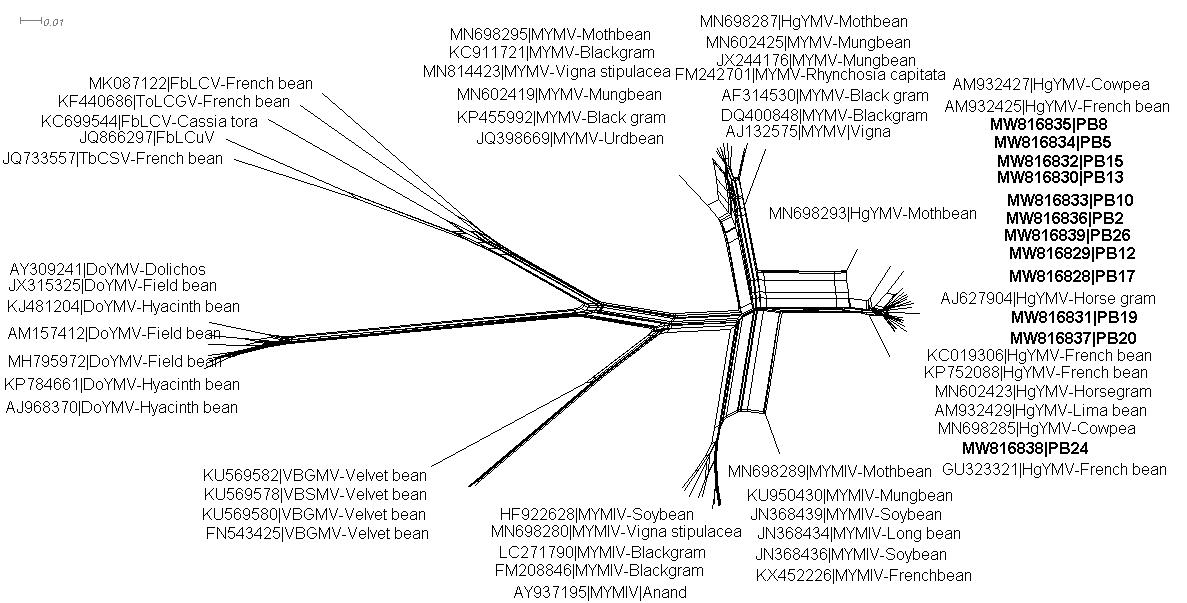

**Figure S4:** A neighbour-net generated for sequences of DNA-A component of 12 HgYMV isolates from pole bean with other selected 50 begomoviruses infecting legumes.

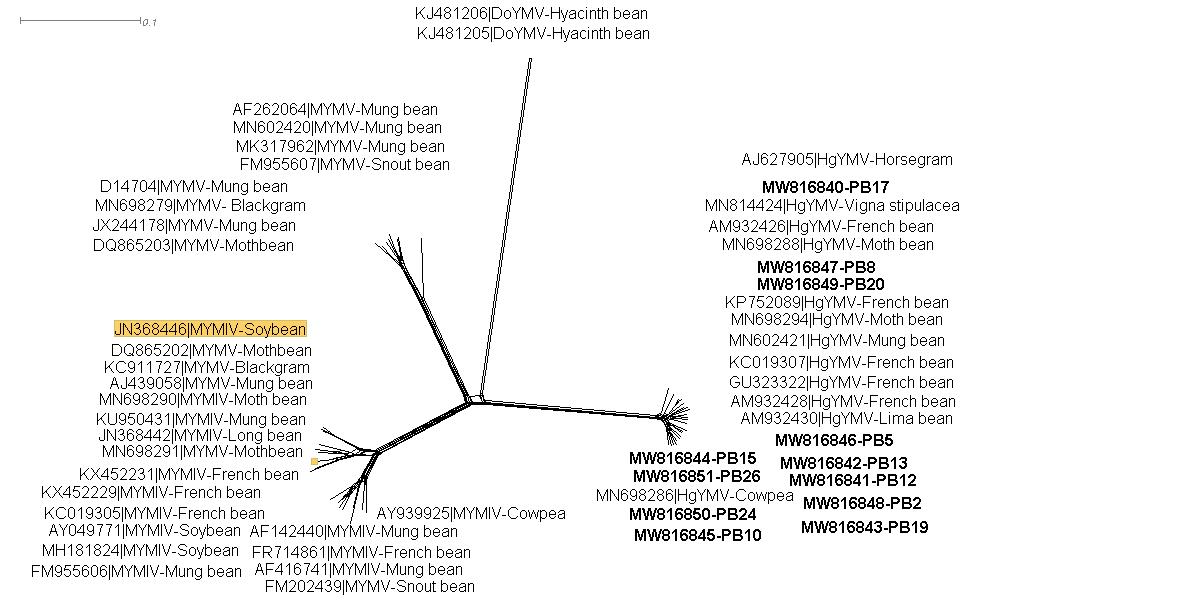

**Figure S5:** A neighbour-net generated for sequences of DNA-B component of 12 HgYMV isolates from pole bean with other selected 42 begomoviruses infecting legumes.

Table S1: List of selected begomoviruses used in the study for comparison analysis of DNA- A component of the 12 pole bean HgYMV isolates

| **Sl.  No.** | **Accession No.** | **Virus** | **Acronym** | **Location** | **Host** |
| --- | --- | --- | --- | --- | --- |
| 1. | MN698285 | *Horsegram yellow mosaic virus* | HgYMV | Belagavi | Cowpea |
| 2. | KC019306 | *Horsegram yellow mosaic virus* | HgYMV | Bengaluru | French bean |
| 3. | KP752088 | *Horsegram yellow mosaic virus* | HgYMV | Bengaluru | French bean |
| 4. | MN698287 | *Horsegram yellow mosaic virus* | HgYMV | Belagavi | Moth bean |
| 5. | MN698293 | *Horsegram yellow mosaic virus* | HgYMV | Belagavi | Moth bean |
| 6. | AM932429 | *Horsegram yellow mosaic virus* | HgYMV | Bengaluru | Lima bean |
| 7. | AM932425 | *Horsegram yellow mosaic virus* | HgYMV | Bengaluru | French bean |
| 8. | MN602423 | *Horsegram yellow mosaic virus* | HgYMV | Dharwad | Horsegram |
| 9. | AM932427 | *Horsegram yellow mosaic virus* | HgYMV | Bengaluru | Cowpea |
| 10. | AJ627904 | *Horsegram yellow mosaic virus* | HgYMV | Coimbatore | Horsegram |
| 11. | GU323321 | *Horsegram yellow mosaic virus* | HgYMV | Sri Lanka | French bean |
| 12. | MN698280 | *Mungbean yellow mosaic India virus* | MYMIV | Raichur | *Vigna stipulacea* |
| 13. | MN698289 | *Mungbean yellow mosaic India virus* | MYMIV | Belagavi | Moth bean |
| 14. | HF922628 | *Mungbean yellow mosaic India virus* | MYMIV | West Bengal | Soybean |
| 15. | KU950430 | *Mungbean yellow mosaic India virus* | MYMIV | Meghalaya | Mung bean |
| 16. | LC271790 | *Mungbean yellow mosaic India virus* | MYMIV | Jabalpur | Blackgram |
| 17. | FM208846 | *Mungbean yellow mosaic India virus* | MYMIV | Pakistan | Blackgram |
| 18. | AY937195 | *Mungbean yellow mosaic India virus* | MYMIV | Anand | - |
| 19. | JN368439 | *Mungbean yellow mosaic India virus* | MYMIV | Indonesia | Soybean |
| 20. | JN368436 | *Mungbean yellow mosaic India virus* | MYMIV | Indonesia | Soybean |
| 21. | JN368434 | *Mungbean yellow mosaic India virus* | MYMIV | Indonesia | Long bean |
| 22. | KX452226 | *Mungbean yellow mosaic India virus* | MYMIV | Oman | French bean |
| 23. | JQ398669 | *Mungbean yellow mosaic virus* | MYMV | New Delhi | Blackgram |
| 24. | MN698295 | *Mungbean yellow mosaic virus* | MYMV | Belagavi | Moth bean |
| 25. | MN602425 | *Mungbean yellow mosaic virus* | MYMV | Dharwad | Mung bean |
| 26. | MN814423 | *Mungbean yellow mosaic virus* | MYMV | Raichur | *Vigna stipulacea* |
| 27. | MN602419 | *Mungbean yellow mosaic virus* | MYMV | Dharwad | Mung bean |
| 28. | KP455992 | *Mungbean yellow mosaic virus* | MYMV | Tirupati | Blackgram |
| 29. | KC911721 | *Mungbean yellow mosaic virus* | MYMV | Coimbatore | Blackgram |
| 30. | JX244176 | *Mungbean yellow mosaic virus* | MYMV | Vietnam | Mung bean |
| 31. | DQ400848 | *Mungbean yellow mosaic virus* | MYMV | Vamban | Blackgram |
| 32. | AJ132575 | *Mungbean yellow mosaic virus* | MYMV | India | Vigna |
| 33. | AF314530 | *Mungbean yellow mosaic virus* | MYMV | Maharashtra | Blackgram |
| 34. | FM242701 | *Mungbean yellow mosaic virus* | MYMV | Pakistan | Snout bean |
| 35. | KJ481204 | *Dolichos yellow mosaic virus* | DoYMV | India | Hyacinth bean |
| 36. | AM157412 | *Dolichos yellow mosaic virus* | DoYMV | Bengaluru | Field bean |
| 37. | KP784661 | *Dolichos yellow mosaic virus* | DoYMV | Tamil Nadu | Hyacinth bean |
| 38. | AJ968370 | *Dolichos yellow mosaic virus* | DoYMV | Mysuru | Hyacinth bean |
| 39. | AY309241 | *Dolichos yellow mosaic virus* | DoYMV | India | Dolichos bean |
| 40. | MH795972 | *Dolichos yellow mosaic virus* | DoYMV | Tamil Nadu | Field bean |
| 41. | JX315325 | *Dolichos yellow mosaic virus* | DoYMV | Bihar | Field bean |
| 42. | KU569578 | *Velvet bean golden mosaic virus* | VbGMV | Taiwan | Velvet bean |
| 43. | KU569580 | *Velvet bean golden mosaic virus* | VbGMV | Taiwan | Velvet bean |
| 44. | KU569582 | *Velvet bean golden mosaic virus* | VbGMV | Taiwan | Velvet bean |
| 45. | FN543425 | *Velvet bean golden mosaic virus* | VbGMV | Taiwan | Velvet bean |
| 46. | JQ866297 | *French bean leaf curl virus* | *FbLCV* | Kanpur | French bean |
| 47. | KF440686 | *Tomato leaf curl Gujarat virus* | ToLCGV | India | French bean |
| 48. | MK087122 | *French bean leaf curl virus* | FbLCV | Madikeri | French bean |
| 49. | KC699544 | *French bean leaf curl virus* | FbLCV | India | Cassia tora |
| 50. | JQ733557 | *Tobacco curly shoot virus* | TbCSV | Varanasi | French bean |

Table S2: List of selected begomoviruses used in the study for comparison analysis of DNA- B component of the 12 pole bean HgYMV s isolates

| **Sl. No.** | **Accession  No.** | **Virus** | **Acronym** | **Location** | **Host** |
| --- | --- | --- | --- | --- | --- |
| 1. | MN698288 | *Horsegram yellow mosaic virus* | HgYMV | Belagavi | Moth bean |
| 2. | MN698286 | *Horsegram yellow mosaic virus* | HgYMV | Belagavi | Cowpea |
| 3. | MN602421 | *Horsegram yellow mosaic virus* | HgYMV | Dharwad | Mung bean |
| 4. | MN814424 | *Horsegram yellow mosaic virus* | HgYMV | Raichur | *Vigna stipulacea* |
| 5. | AM932426 | *Horsegram yellow mosaic virus* | HgYMV | Bengaluru | French bean |
| 6. | AJ627905 | *Horsegram yellow mosaic virus* | HgYMV | Coimbatore | Horsegram |
| 7. | AM932428 | *Horsegram yellow mosaic virus* | HgYMV | Bengaluru | French bean |
| 8. | MN698294 | *Horsegram yellow mosaic virus* | HgYMV | Belagavi | Moth bean |
| 9. | KP752089 | *Horsegram yellow mosaic virus* | HgYMV | Bengaluru | French bean |
| 10. | GU323322 | *Horsegram yellow mosaic virus* | HgYMV | Sri Lanka | French bean |
| 11. | KC019307 | *Horsegram yellow mosaic virus* | HgYMV | Dharwad | French bean |
| 12. | AM932430 | *Horsegram yellow mosaic virus* | HgYMV | Bengaluru | Lima bean |
| 13. | KU950431 | *Mungbean yellow mosaic India virus* | MYMIV | Meghalaya | Mung bean |
| 14. | KX452229 | *Mungbean yellow mosaic India virus* | MYMIV | Oman | French bean |
| 15. | LC271791 | *Mungbean yellow mosaic India virus* | MYMIV | Jabalpur | Blackgram |
| 16. | JN368446 | *Mungbean yellow mosaic India virus* | MYMIV | Indonesia | Soybean |
| 17. | JN368442 | *Mungbean yellow mosaic India virus* | MYMIV | Indonesia | Long bean |
| 18. | KX452231 | *Mungbean yellow mosaic India virus* | MYMIV | Oman | French bean |
| 19. | FR714861 | *Mungbean yellow mosaic India virus* | MYMIV | Palampur | French bean |
| 20. | MH181824 | *Mungbean yellow mosaic India virus* | MYMIV | India | Soybean |
| 21. | AF416741 | *Mungbean yellow mosaic India virus* | MYMIV | India | Mung bean |
| 22. | MN698290 | *Mungbean yellow mosaic India virus* | MYMIV | Belagavi | Moth bean |
| 23. | KC019305 | *Mungbean yellow mosaic India virus* | MYMIV | Varanasi | French bean |
| 24. | FM955606 | *Mungbean yellow mosaic India virus* | MYMIV | Pakistan | Mung bean |
| 25. | AY939925 | *Mungbean yellow mosaic India virus* | MYMIV | New Delhi | Cowpea |
| 26. | FM202439 | *Mungbean yellow mosaic India virus* | MYMIV | Pakistan | Snout bean |
| 27. | AY049771 | *Mungbean yellow mosaic India virus* | MYMIV | India | Soybean |
| 28. | AF142440 | *Mungbean yellow mosaic India virus* | MYMIV | India | Mung bean |
| 29. | MN698279 | *Mungbean yellow mosaic virus* | MYMV | Raichur | Blackgram |
| 30. | JX244178 | *Mungbean yellow mosaic virus* | MYMV | Vietnam | Mung bean |
| 31. | D14704 | *Mungbean yellow mosaic virus* | MYMV | India | Mung bean |
| 32. | AF262064 | *Mungbean yellow mosaic virus* | MYMV | India | Mung bean |
| 33. | MK317962 | *Mungbean yellow mosaic virus* | MYMV | Coimbatore | Mung bean |
| 34. | MN602420 | *Mungbean yellow mosaic virus* | MYMV | Dharwad | Mung bean |
| 35. | DQ865203 | *Mungbean yellow mosaic virus* | MYMV | Tamil Nadu | Mothbean |
| 36. | FM955607 | *Mungbean yellow mosaic virus* | MYMV | Pakistan | Snout bean |
| 37. | MN698291 | *Mungbean yellow mosaic virus* | MYMV | Belagavi | Moth bean |
| 38. | DQ865202 | *Mungbean yellow mosaic virus* | MYMV | Tamil Nadu | Mothbean |
| 39. | AJ439058 | *Mungbean yellow mosaic virus* | MYMV | Dharwad | Mung bean |
| 40. | KC911727 | *Mungbean yellow mosaic virus* | MYMV | Vamban | Blackgram |
| 41. | KJ481205 | *Dolichos yellow mosaic virus* | DoYMV | India | Hyacinth bean |
| 42. | KJ481206 | *Dolichos yellow mosaic virus* | DoYMV | India | Hyacinth bean |

**Table 1:** Percent disease incidence of yellow mosaic disease of pole bean in Kolar, Chikkaballapura and Bengaluru Rural districts during 2019-2020

| **District** | **Taluk** | **Village** | **Longitude** | **Latitude** | **Symptoms observed** | **Variety grown** | **Isolate** | **PDI (%)** | **PDI of taluk (%)** | **PDI of district (%)** |
| --- | --- | --- | --- | --- | --- | --- | --- | --- | --- | --- |
| Kolar | Malur | Agrahara | 13.3141 ^0^N | 78.1529 ^0^E | Vein clearing, mosaic, yellowing | Blue belli | PB1 | 10.19 | 21.63 | 33.91 |
|  |  | Anepura | 13.0337 ^0^N | 78.0427 ^0^E | Mosaic, vein clearing | Blue belli | PB2 | 40.12 |  |  |
|  |  | Chikkakunthur | 13.0033 ^0^N | 78.0136 ^0^E | Vein clearing, mosaic, yellowing | Blue belli | PB3 | 20.13 |  |  |
|  |  | Hungenahalli | 13.0421 ^0^N | 77.9921 ^0^E | Mosaic, yellowing | Blue belli | PB4 | 15.18 |  |  |
|  | Kolar | Balagere | 13.2987 ^0^N | 78.0902 ^0^E | Mosaic, yellowing | Blue belli | PB5 | 30.12 | 27.58 |  |
|  |  | Maduvathii | 13.0695 ^0^N | 78.0888 ^0^E | Mosaic, yellowing | Blue belli | PB6 | 25.05 |  |  |
|  | Bangarpet | Nayakarapalli | 13.0528 ^0^N | 78.2038 ^0^E | Mosaic, yellowing | Blue belli | PB7 | 6.020 | 10.95 |  |
|  |  | Siddenahalli | 13.2213 ^0^N | 77.9100 ^0^E | Vein clearing, mosaic, yellowing | Blue belli | PB8 | 15.82 |  |  |
|  | Mulbagal | Chiteri | 13.1667 ^0^N | 78.2400 ^0^E | Mosaic, yellowing | Blue belli | PB9 | 29.02 | 33.72 |  |
|  |  | Doddiganahalli | 13.1944 ^0^N | 77.9382 ^0^E | Mosaic, yellowing | Blue belli | PB10 | 38.43 |  |  |
|  | Srinivaspura | Pathapalli | 13.2016 ^0^N | 78.1243 ^0^E | Mosaic, yellowing, deformed fruit | NZ super King | PB11 | 70.61 | 75.68 |  |
|  |  | Gundamnatta | 13.3418 ^0^N | 78.2132 ^0^E | Mosaic, yellowing | Blue belli | PB12 | 80.74 |  |  |
| Chikkaballapur | Chintamani | Nandiganahalli | 13.5659 ^0^N | 77.5318 ^0^E | Mosaic, yellowing | NZ super King | PB13 | 72.93 | 48.18 | 33.29 |
|  |  | Chokkanahalli | 13.4360 ^0^N | 78.0276 ^0^E | Vein clearing, mosaic, yellowing | NZ super King | PB14 | 23.43 |  |  |
|  | Chikkaballapur | Nandhi | 13.2150 ^0^N | 77.4101 ^0^E | Mosaic, yellowing | Blue belli | PB15 | 35.17 | 20.97 |  |
|  |  | Malemachanahalli | 13.3410 ^0^N | 77.8619 ^0^E | Mosaic, yellowing | Blue belli | PB16 | 06.78 |  |  |
|  | Sidlaghatta | Suguturu | 13.3914 ^0^N | 77.8649 ^0^E | Mosaic, yellowing | Blue belli | PB17 | 40.02 | 36.10 |  |
|  |  | Ankathatti | 13.3049 ^0^N | 77.8181 ^0^E | Mosaic, yellowing | Blue belli | PB18 | 32.18 |  |  |
|  | Bagepalli | Kammaravaripalli | 13.7800 ^0^N | 77.7900 ^0^E | Vein clearing, mosaic, yellowing | NZ super King | PB19 | 27.92 | 27.92 |  |
| Bangalore Rural | Nelamangala | K.G Lekkanahall | 13.0185 ^0^N | 77.4573 ^0^E | Mosaic, yellowing | NZ super King | PB20 | 28.23 | 20.73 | 28.77 |
|  |  | Hasurahalli | 13.1925 ^0^N | 77.3652 ^0^E | Mosaic, vein clearing | NZ super King | PB21 | 18.18 |  |  |
|  |  | Narasapura | 13.0998 ^0^N | 77.3936 ^0^E | Mosaic, yellowing | Blue belli | PB22 | 15.14 |  |  |
|  |  | Thyamagondlu | 13.2142 ^0^N | 77.2994 ^0^E | Mosaic, yellowing | Blue belli | PB23 | 21.37 |  |  |
|  | Hoskote | Sulibele | 13.1886 ^0^N | 77.7881 ^0^E | Mosaic, yellowing, deformed fruit | Blue belli | PB24 | 24.03 | 18.15 |  |
|  |  | Allappanahalli | 13.0810 ^0^N | 77.7886 ^0^E | Vein clearing, mosaic, yellowing | Blue belli | PB25 | 12.27 |  |  |
|  | Doddaballapura | Hadonahalli | 13.29875 ^0^N | 77.5409 ^0^E | Mosaic, yellowing | NZ super King | PB26 | 47.34 | 47.34 |  |

| **Begomoviruses** | **DNA A** | **Intergenic  region (IR)** |  | **Replication  initiation protein (AC1)** | **Transcriptional activator  protein (AC2)** | **Replication enhancer protein (AC3)** | **C4 (AC4)** | **Coat protein  (AV1)** | **Pre-coat  protein (AV2)** |
| --- | --- | --- | --- | --- | --- | --- | --- | --- | --- |
| **Within the  current 12 isolates** | 95.10-98.20 | 85.40-99.20 | **nt** | 95.50-100.00 | 90.40-100.00 | 90.60-98.00 | 96.90-100.00 | 90.50-100.00 | 95.40-100.00 |
|  |  |  | **aa** | 93.90-100.00 | 86.10-100.00 | 82.80-99.20 | 93.80-100.00 | 85.40.100.00 | 92.80-100.00 |
| **HgYMV *(11)** | 92.70-98.00 | 40.10-99.60 | **nt** | 95.60-98.50 | 96.00-99.50 | 90.10-99.00 | 96.50-99.60 | 82.90-95.80 | 93.40-99.40 |
|  |  |  | **aa** | 96.10-98.60 | 88.10-99.20 | 66.10-82.80 | 91.70-98.90 | 78.70-93.30 | 88.30-98.20 |
| **MYMIV *(11)** | 77.60-86.50 | 31.50-77.70 | **nt** | 83.10-85.00 | 85.70-87.90 | 84.10-86.10 | 81.00-83.30 | 71.70-93.60 | 77.20-86.30 |
|  |  |  | **aa** | 83.10-85.90 | 77.00-82.20 | 76.80-97.00 | 60.00-68.60 | 56.20-86.70 | 69.60-7190 |
| **MYMV *(12)** | 82.20.-84.70 | 29.00-72.90 | **nt** | 83.70-85.70 | 88.70-92.80 | 81.70-95.8 | 85.70-89.10 | 74.10-79.50 | 79.20-83.70 |
|  |  |  | **aa** | 78.70-87.20 | 82.30-89.60 | 58.20-64.90 | 70.10-78.30 | 54.20-66.90 | 66.30-73.40 |
| **VbGMV *(4)** | 73.40 -74.90 | 52.80-58.70 | **nt** | 69.50-74.10 | 76.40-79.90 | 70.80-76.00 | 50.40-51.80 | 70.70-75.20 | 67.50-71.60 |
|  |  |  | **aa** | 69.00-74.00 | 63.20-68.10 | 55.10-59.50 | 55.60-60.20 | 52.50-56.40 | 55.50-60.00 |
| **DoYMV *(7)** | 61.70-62.70 | 33.90-49.60 | **nt** | 60.00-61.70 | 62.60-64.90 | 38.90-64.10 | 63.20-69.90 | 63.40-67.60 | 47.20-50.10 |
|  |  |  | **aa** | 58.10-60.00 | 51.40-57.00 | 55.10-59.50 | 54.40-58.00 | 40.60-46.60 | 12.80-31.30 |
| **FbLCV**  ***(3)** | 62.30-63.50 | 30.50-47.80 | **nt** | 34.10-69.60 | 53.80-60.90 | 53.50-56.30 | 63.20-69.90 | 60.30-64.20 | 54.10-58.90 |
|  |  |  | **aa** | 12.30-73.20 | 36.10-48.10 | 38.00-40.20 | 38.80-41.20 | 40.80-46.60 | 36.10-56.40 |
| **ToLCGV *(1)** | 63.50-64.00 | 36.00-55.10 | **nt** | 71.7-72.50 | 58.50-59.50 | 57.50-58.30 | 77.20-78.20 | 58.50-61.90 | 58.60-60.10 |
|  |  |  | **aa** | 70.40-71.20 | 42.90-45.10 | 58.60-61.30 | 57.70-61.80 | 36.40-41.100 | 36.50-38.20 |
| **TbCSV *(1)** | 63.80-64.40 | 35.30-38.70 | **nt** | 70.50-71.40 | 58.70-60.20 | 56.90-58.10 | 75.80-77.20 | 58.40-62.70 | 55.80-57.80 |
|  |  |  | **aa** | 68.70-69.80 | 40.70-47.40 | 57.70-64.10 | 56.70-60.80 | 37.00-40.90 | 36.20-37.90 |

**Table-2:** Nucleotide and amino acid identity of DNA A of 12 HgYMV isolates from pole beanwith other selected 42 begomoviruses infecting legume

**Table 3**: Nucleotide and amino acid identity of DNA B of 12 HgYMV isolates from pole bean with other selected 42 begomoviruses infecting legume

| **Begomoviruses** | **DNA B** | **Intergenic region (IR)** | **Movement protein (BC1)** | | **Nuclear shuttle protein (BV1)** |
| --- | --- | --- | --- | --- | --- |
| **Within the current 12 isolates** | 94.70-97.40 | 93.40-98.50 | **nt** | 97.90-100.00 | 94.40-98.90 |
|  |  |  | **aa** | 95.79-100.00 | 89.00-98.80 |
| **HgYMV *(12)** | 93.70 - 97.40 | 92.30 - 99.10 | **nt** | 95.40 - 99.10 | 94.20 - 99.20 |
|  |  |  | **aa** | 77.80 - 100.00 | 91.00 - 98.40 |
| **MYMIV *(15)** | 61.60 - 68.90 | 43.70 - 47.50 | **nt** | 77.10 - 80.70 | 77.80 - 81.70 |
|  |  |  | **aa** | 19.70 - 86.50 | 68.70 - 73.00 |
| **MYMV *(13)** | 62.10 - 68.30 | 42.50 - 51.70 | **nt** | 61.20 - 80.60 | 64.30 - 81.80 |
|  |  |  | **aa** | 83.50 - 87.20 | 57.40 - 73.00 |
| **DoYMV *(2)** | 52.10 – 52.60 | 41.10 - 41.30 | **nt** | 64.90 - 66.40 | 60.50 - 67.70 |
|  |  |  | **aa** | 69.70 - 71.00 | 48.60 - 51.30 |
